# MKS-NPHP module proteins regulate ciliary shedding in *Paramecium*

**DOI:** 10.1101/676395

**Authors:** Delphine Gogendeau, Michel Lemullois, Anne Aubusson-Fleury, Olivier Arnaiz, Jean Cohen, Christine Vesque, Sylvie Schneider-Maunoury, France Koll, Anne-Marie Tassin

**Affiliations:** Institute for Integrative Biology of the Cell (I2BC), CEA, CNRS, Univ. Paris Sud, Université Paris-Saclay, 1 Avenue de la Terrasse, 91198 Gif sur Yvette, France; Sorbonne Université, CNRS UMR7622, INSERM U1156, Developmental Biology Laboratory - Institut de Biologie Paris-Seine (IBPS), Paris, France

## Abstract

Ciliogenesis is a general process in eukaryotic cells and its different steps begin to be well characterised. However, the molecular mechanisms leading to decilation or ciliary shedding are still poorly understood. This process, observed from unicellular organisms such as *Chlamydomonas* or *Paramecium* to multiciliated cells from trachea or fallopian tube of vertebrates, seems to be a general process since recent observations demonstrates its requirement during the cell cycle or neurogenesis. Interestingly, in all cellular models, ciliary shedding occurs distal to the transition zone, essentially known to act as a diffusion barrier between the intracellular space and the cilium, suggesting conserved molecular mechanisms.

To determine if MKS and NPHP modules, known to cooperate to establish transition zone formation and function, could control ciliary shedding, we studied in *Paramecium* the function of TMEM216/MKS2 and TMEM107 (two members of the MKS module), NPHP4 (one member of the NPHP module), CEP290/NPHP6 and RPGRIP1L/MKS5. We show that all these proteins are recruited to the TZ as soon as growing cilia are detected and localise with a 9-fold symmetry at the level of the axonemal plate. Interestingly, we demonstrate that the depletion of the two MKS module proteins induces spontaneous cilia shedding, while the depletion of either NPHP4, CEP290 or RPGRIP1L inhibits the process. Our results constitute the first evidence for a role of conserved TZ proteins in deciliation and open new directions for understanding motile cilia physiology.

## INTRODUCTION

Cilia are highly conserved cell appendages endowed with motility and sensory functions acting as “cell antennae”. Primary cilia that emanate from the surface of most cells in multicellular organisms are key organelles in numerous developmental and physiological processes (Goetz & Anderson, 2010). Cilia conserved architecture consists of three structural regions: the basal body (BB) at their proximal part, the axoneme composed of nine microtubule doublets covered by the ciliary membrane at the distal part and in between, the transition zone (TZ). The transition zone is characterised by Y-shaped linkers connecting the microtubule doublets of the axoneme to the ciliary membrane, the so-called Y-links, and the transition fibers, which are thought to function together as a ciliary gate, ensuring a specific ciliary composition. Genetic studies together with protein-protein interaction data and proteomics analyses, led to the identification of various components of this TZ (Williams *et al*, 2011, Huang *et al*, 2011, Garcia-Gonzalo *et al*, 2011, Sang *et al*, 2011, Diener *et al*, 2015, Dean *et al*, 2016, for review see Reiter *et al*, 2012, Gonçalves & Pelletier, 2017). Some of them are assembled in two distinct protein complexes, defined as the MKS and the NPHP modules, which cooperate in biogenesis and function of the transition zone (Williams *et al*, 2011). By recruiting the proteins of these complexes, CEP290 and RPGRIP1L, play a crucial function in TZ assembly (Li *et al*, 2016, Schouteden *et al*, 2015, Basiri *et al*, 2014, Wiegering *et al*, 2018b). These proteins are supposed to be associated with the Y-links (Craige *et al*, 2010). Importantly, many TZ proteins encoding genes are mutated in human ciliopathies (Czarnecki & Shah, 2012, Mitchison & Valente, 2017)

Ciliogenesis is a multi-step process, including duplication and maturation of the centriole/basal body, followed by its anchoring to ciliary vesicle membranes inside the cell or to the cell surface via the plasma membrane. Finally, BB templates axonemal growth thanks to intraflagellar transport.

While the ciliogenesis process is now well documented, the deciliation process (also called flagellar excision or autotomy) is still poorly understood. Deciliation is different from cilia resorption/involution, and is characterised by the shedding of cilia or flagella above the membrane at the distal end of the transition zone, often in response to a stress, (Satir *et al*, 1976, Adoutte *et al*, 1980b, Quarmby, 2004a). Deciliation is widely used in unicellular organisms such as *Chlamydomonas*, *Paramecium* or *Tetrahymena,* but also occurs in vertebrates in multiciliated epithelia such as the upper airway or oviduct. Physiological deciliation of the oviduct, has been observed both in birds and in mammals during the luteal phase of the menstrual cycle (Brenner, 1969, Verhage *et al*, 1979, Odor *et al*, 1980, Boisvieux-Ulrich *et al*, 1980). This process is more pronounced after progesterone-therapy (Donnez *et al*, 1985). Smoke and nitrogen dioxide inhalation (Heller & Gordon, 1986) were shown to induce deciliation in the upper airway, the breakage occurring just above the ciliary necklace. During upper airway infection by *Mycoplasma pneumonia or Bordetella pertussis,* the disorganisation of the ciliary necklace, and in some cases its absence, have been observed (Carson *et al*, 1980, Muse *et al*, 1977, Wilson *et al*, 1991), suggesting a strong effect of bacterial infection on the transition zone, which could lead to subsequent loss of cilia.

In *Chlamydomonas,* deflagellation is triggered by an increase in intracellular Calcium near the base of the flagella (Quarmby & Hartzell, 1994, Quarmby, 1996 Wheeler *et al*, 2008) and involves microtubule severing proteins such as katanin (Lohret *et al*, 1998) and FA1p, a non evolutionarily conserved protein (Finst *et al*, 1998, Finst *et al*, 2000). Interestingly, proteins involved in this process have been shown to localise at the TZ (Finst *et al*, 2000, Mahjoub *et al*, 2002).

In order to evaluate a possible involvement of the conserved MKS/NPHP TZ proteins in the deciliation process of motile cilia, we studied five proteins belonging to these modules including CEP290 and RPGRIP1L in *Paramecium,* a free-living unicellular organism that bears at its surface ca. 4000 cilia. We show using super-resolution microscopy and TEM that all five proteins localise with nine fold symmetry at the distal part of the TZ, precisely at the level of the axosomal plate, which is consistent with their localisation both in non-motile and motile cilia of various organisms (Jana *et al*, 2018, Gonçalves & Pelletier, 2017). Functional studies using RNAi showed that, as expected, the depletion of TZ proteins impairs TZ assembly and function. Most interestingly, we demonstrated that the depletion of any of these proteins impacts the deciliation process. While the depletion of the two MKS protein modules induces spontaneous cilia autotomy, the depletion of either NPHP4, CEP290 or RPGRIP1L increases resistance to deciliation induced by Ca^2+^- ethanol treatment. Our results constitute the first evidence for a role of these TZ proteins in deciliation and open new directions for understanding cilia physiology.

## RESULTS

To assess the involvement of conserved TZ proteins in ciliary shedding, bio-informatics analyses were undertaken and showed that CEP290, RPGRIP1L as well as most MKS and NPHP module proteins are conserved in *Paramecium* (Table 1, our personal analyses, Barker *et al*, 2014, Schouteden *et al*, 2015). We focused on five of them: two belonging to the MKS module (TMEM216/MKS2 and TMEM107), one representative of the NPHP complex (NPHP4) and the two pivotal proteins, CEP290/NPHP6 and RPGRIP1L/MKS5/NPHP8. To simplify, we will refer to them as a whole as “TZ proteins”. Due to whole genome duplications, which occurred during evolution in this lineage (Aury *et al*, 2006), several ohnologs (paralogs issued from whole genome duplication) (Ohno S., 1970) of each protein are encoded in the *Paramecium tetraurelia* genome (See Table S1).

**Table 1:**
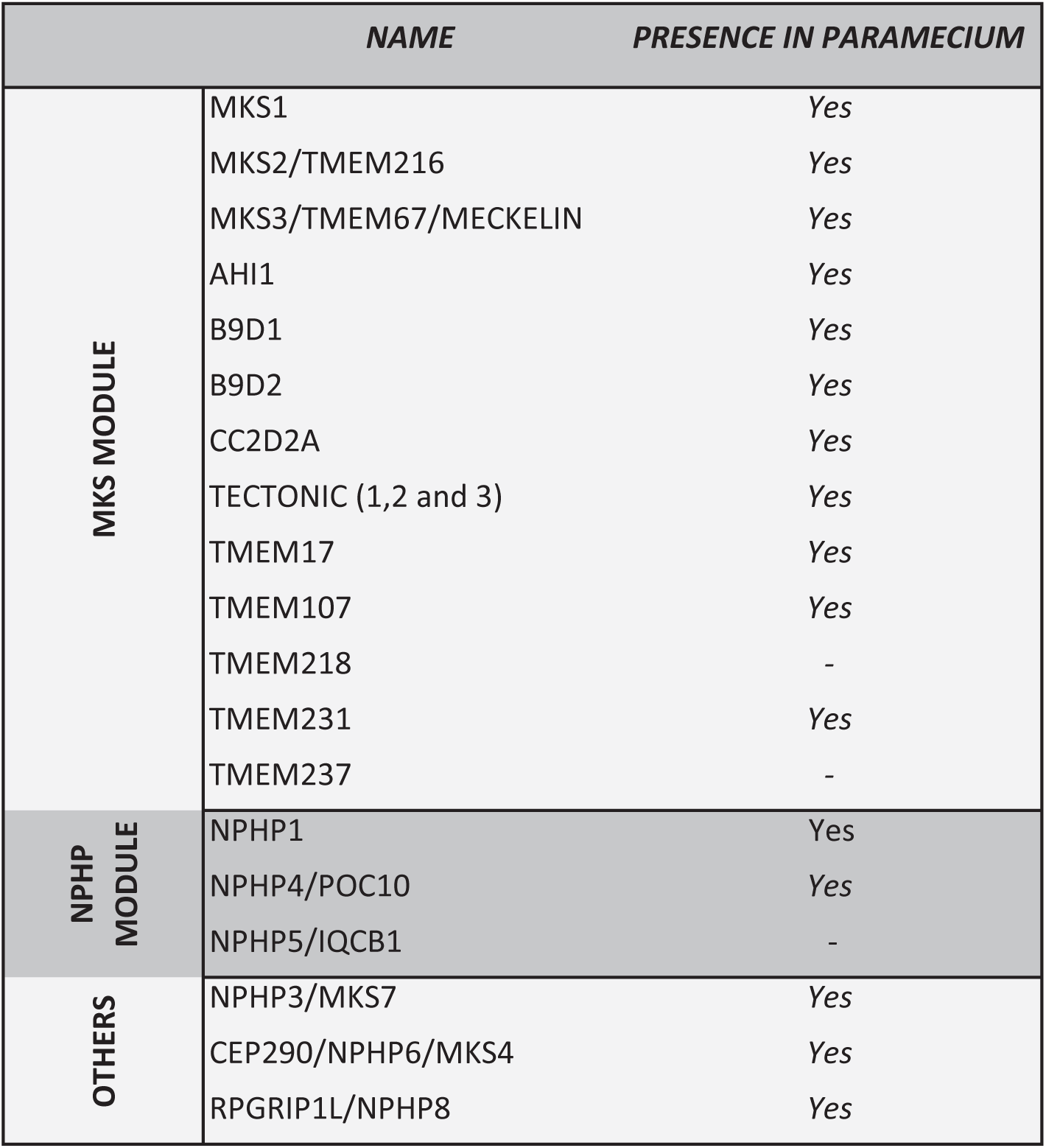
Conservation of transition zone proteins in *Paramecium*.

| NAME |  | PRESENCE IN PARAMECIUM |
| --- | --- | --- |
| MKS MODULE | MKS1 | Yes |
|  | MKS2/TMEM216 | Yes |
|  | MKS3/TMEM67/MECKELIN | Yes |
|  | AHI1 | Yes |
|  | B9D1 | Yes |
|  | B9D2 | Yes |
|  | CC2D2A | Yes |
|  | TECTONIC (1,2 and 3) | Yes |
|  | TMEM17 | Yes |
|  | TMEM107 | Yes |
|  | TMEM218 | - |
|  | TMEM231 | Yes |
|  | TMEM237 | - |
| NPHP MODULE | NPHP1 | Yes |
|  | NPHP4/POC10 | Yes |
|  | NPHP5/IQCB1 | - |
| OTHERS | NPHP3/MKS7 | Yes |
|  | CEP290/NPHP6/MKS4 | Yes |
|  | RPGRIP1L/NPHP8 | Yes |

### TZ proteins are recruited at the axosomal plate at the onset of ciliogenesis

In a first set of experiments, we assessed whether the candidate proteins localised at the transition zone like their counterparts in other organisms. This was performed by expressing one GFP-tagged ohnolog of each protein.

Paramecia display a precise BB and cilia organization (Iftode *et al*, 1989, Aubusson-Fleury *et al*, 2012, for review see Tassin *et al*, 2015, Figure S1). In the anterior part of the cell, the invariant field shows cortical units with mostly 2 ciliated BBs. The posterior part of the cell displays units with a single BB, and the mixed field exhibit units with either a single or 2 BBs. In the latter case, both BBs are anchored at the cell surface but only the posterior one is ciliated (Figure S1). Units with a single BB can be ciliated or not according to the cell cycle stage. In *Paramecium,* the TZ displays three plates, from proximal to distal: the terminal plate, the intermediate plate and the axosomal plate. Interestingly, ultrastructural analysis of the TZ of unciliated or ciliated BBs reveals that they have a distinct morphology, the ciliated ones showing a longer TZ, suggesting a structural maturation of the TZ during the ciliation process. (Figure S1, Aubusson-Fleury *et al*, 2012, Tassin *et al*, 2015). After saponin extraction and fixation, microscopic analysis performed on each transformed cell line, detected GFP-fluorescence at the transition zone of ciliated BBs as demonstrated in Figure 1A and supplemental Figure S2A and interestingly was not detected at the TZ of unciliated BBs. These results suggest that molecular maturation of the TZ accompanies the structural maturation.

**Figure 1:**
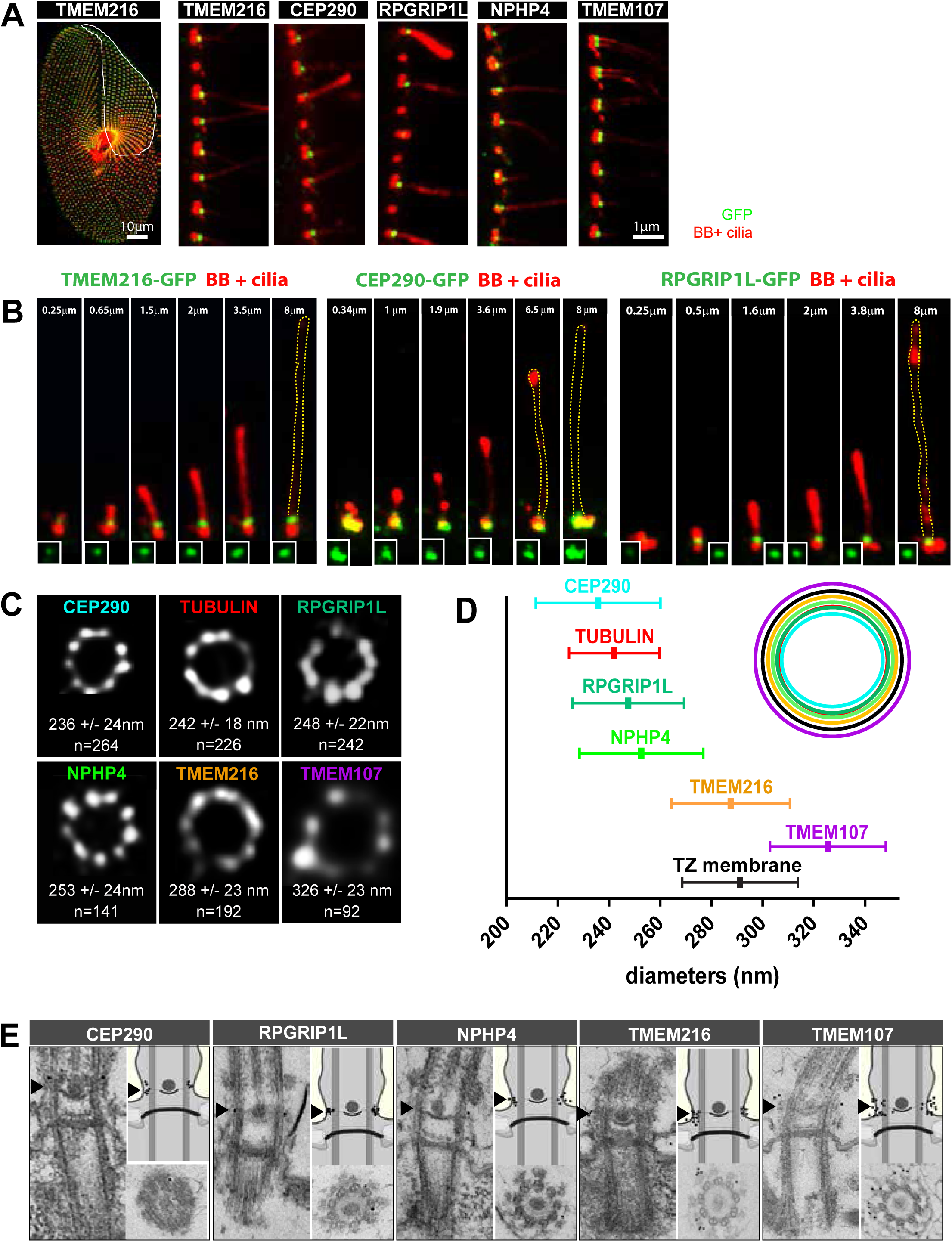
Localisation of *Paramecium* Transition zone proteins. A- Paramecia expressing different TZ proteins fused with GFP. Cells were permeabilized before proceeding to immunostaining. Left panels: surface view of a paramecium double stained by the monoclonal anti-glutamylated tubulin ID5 (decorating basal bodies (BB) and cilia in red) and a polyclonal anti-GFP (in green). Ciliated basal bodies of the invariant field (encircled in white) are stained by both GFP antibodies and ID5. In the other part of the cells some basal bodies appeared only labelled by ID5 and represented unciliated basal bodies. Right panels: optical sections, transverse to the cell surface at the cell margin showing a ciliary row double labelled by the polyclonal ID5 antibodies (decorating BB and cilia) and an anti-GFP antibodies (green). Note that ID5 antibodies better recognise short cilia. TMEM107-GFP, RPGRIP1L-GFP and TMEM216-GFP are exclusively localised at the transition zone of ciliated basal bodies. CEP290-GFP and NPHP4-GFP in addition to their localisation at the TZ of ciliated basal bodies can also be observed on basal bodies BB. Bars =10μm and 1μm. B- Cilia at different steps of their growth labelled by ID5, which decorated also the basal body. To ascertain that cilia were growing and not broken, dividing cells where new cilia are generated were observed. To avoid fluctuations in fluorescence intensity due to variations in expression rate, all the cilia were extracted within a same cell for each expressing TZ-GFP as indicated. Using an anti-GFP antibodies, the GFP signal is detected at the TZ as soon as the growing cilium is detected by ID5 antibodies. C- Representative STED images revealing distinct localisation patterns of several GFP-tagged TZ proteins labelled with anti-GFP or ID5 (tubulin). A single ring differing in diameter is observed according to the observed protein. Measurements of the mean diameters (distances between intensity maxima) and the number of BB analysed are given beneath each image. D- For quantification, distances between intensity maxima of the toroid were measured and plotted as the mean+/-sd (see C) for each GFP-tagged TZ protein. A ring of the color corresponding to the protein and representing the localisation of each protein is shown on the right. E- Representative EM images of the immuno-localisation of the different GFP fusion labelling revealed by an anti-GFP antibodies. Left panels: longitudinal views. Lower right panels: Transverse section of a BB at the level of the axosomal plate. Upper right panels: BB scheme recapitulating the localisation of gold beads. All these proteins, although occupying different diameters in transverse views, are localised at the level of the axosomal plate indicated by a black arrowhead. CEP290: 11 gold beads on 10 BB; RPGRIP1L: 24 gold beads on 18 BB; NPHP4: 14 gold beads on 8 BB; TMEM107: 35 gold beads on 16 BB; TMEM216: 36 gold beads on 21 BB.

In addition to a common localisation at the TZ, NPHP4-GFP and CEP290-GFP are also present on the proximal part of all BBs (Figure 1A). An additional cortical staining associated with all basal bodies was also observed for TMEM216-GFP on living or fixed cells without detergent extraction (Figure S2A) suggesting that two pools of TMEM216 coexist in the cell: a soluble pool associated with all BBs and an insoluble pool, restricted to the TZ of ciliated BBs. In the subsequent analyses, we focused only on the TZ labelling of each of the five proteins.

The localisation of all studied proteins in the TZ of ciliated BBs led us to study the recruitment of these proteins during ciliary growth. This analysis demonstrated that each protein was recruited at the TZ as soon as the growing cilium is detected by the antibodies (Figure 1B and supplemental Figure 2A). We then combined super-resolution microscopy (STED) and immuno-electron microscopy observations to increase localisation accuracy (Figures 1C and D). All proteins were organized into a nine-fold symmetry. Quantification of the mean diameters of the rings revealed that CEP290-GFP localised most centrally, close to the microtubules (236 +/-24nm, n=264 for CEP290 vs 242+/-18 nm, n=226 for tubulin), while both RPGRIP1L-GFP (247 +/-22nm, n=242) and NPHP4-GFP (253+/-24 nm, n=141) were detected outside the ring of microtubule doublets. Finally, TMEM216-GFP and TMEM107-GFP (diameters 288 +/- 23nm, n=192 and 326 +/-23nm, n=92 respectively) were found more externally (Figure 1C), close to the membrane positioned on the graph Figure 1D. Immuno-EM data confirmed these observations and suggested that TMEM216-GFP as well as TMEM107-GFP are closely localised to the TZ ciliary membrane. Interestingly, a clear restriction of all these proteins to the axosomal plate (arrowheads on Figure 1E) is found. This localisation in *Paramecium* corresponds to the localisation of Y-links, the connection of transition fibers to the ciliary membrane and the ciliary necklace (Dute & Kung, 1978).

### Depletion of TZ proteins affects proliferation rate and swimming velocities

To ascertain the function of these TZ proteins, all ohnologs of each protein family were depleted concurrently by inactivating all the corresponding genes (TZ^RNAi^) using the feeding method (Galvani & Sperling, 2002, Carradec *et al*, 2015). The efficiency of silencing was tested by controlling the effective depletion of the GFP-tagged protein (Supplemental Figure S3). Within the first 24h, depleted cells showed an abnormal swimming behaviour associated with a slow growth rate (1 division less than controls per 24h, data not shown). These phenotypes were stably maintained for several days in the RNAi media except for TMEM216/MKS2, which died after 72h.

To better characterise the motility defects in silenced cells, we video-recorded paramecia (Supplemental movies 1-6). Control cells had an average swimming velocity of 683 μm per second (Figure S4, n=214 or n=232 cells, for 24h and 48h respectively). Depletion of TZ proteins resulted in a severe reduction (by about 1/3) of this velocity (Figure S3) similar to the one observed after depletion of Bug22 (Laligné *et al*, 2010), and of DNAH9 or C11orf70, these last two proteins being required for dynein arms assembly (Fassad *et al*, 2018a, b). For all our subsequent analyses, we used cells cultured for 48h under the RNAi conditions (unless otherwise stated).

### Depletion of TMEM107 and TMEM216 leads to an accumulation of stretches of short cilia

To determine whether the swimming alterations could be due to defects in ciliation or in BB organisation, the ciliation patterns of the RNAi-treated cells were analysed by immunolabelling. To avoid possible artefacts in the labelling, control RNAi and TZ^RNAi^ cells were mixed and processed together in all IF experiments (see Materials and Methods). In all silenced cells, the BB pattern was normal (Supplemental Figure 5) indicating that TZ protein depletion did not affect BB duplication or anchoring processes. During interphase, control paramecia mainly exhibit long fully assembled cilia with sparse developing cilia dispersed all along the same ciliary row. Cep290^RNAi^, NPHP4^RNAi^ and RPGRIP1L^RNAi^ cells displayed a ciliary pattern essentially identical to the control cells (Figure 2A). In contrast, interphasic TMEM107^RNAi^ or TMEM216^RNAi^ cells displayed an accumulation of stretches of short cilia, measuring less than 5 μm (Figure 2B). Such short cilia were never observed in control cells or after depletion of either CEP290, NPHP4 or RPGRIP1L as observed on optical sections, transverse to the cell surface at the cell margin. Quantification of these short cilia (Figure 2C) confirmed a dramatic increase in their number for both TMEM107^RNAi^ (25.8 +/- 2%, N=50 cells) and TMEM216^RNAi^ (26.55 +/- 1.7%, N=42 cells), compared to control cells (mean percentage of 9.92 +/- 0.5%, N=60 cells). Several hypotheses may explain this phenotype such as slow ciliary growth, ciliary involution or constant deciliation followed by ciliary regrowth.

**Figure 2:**
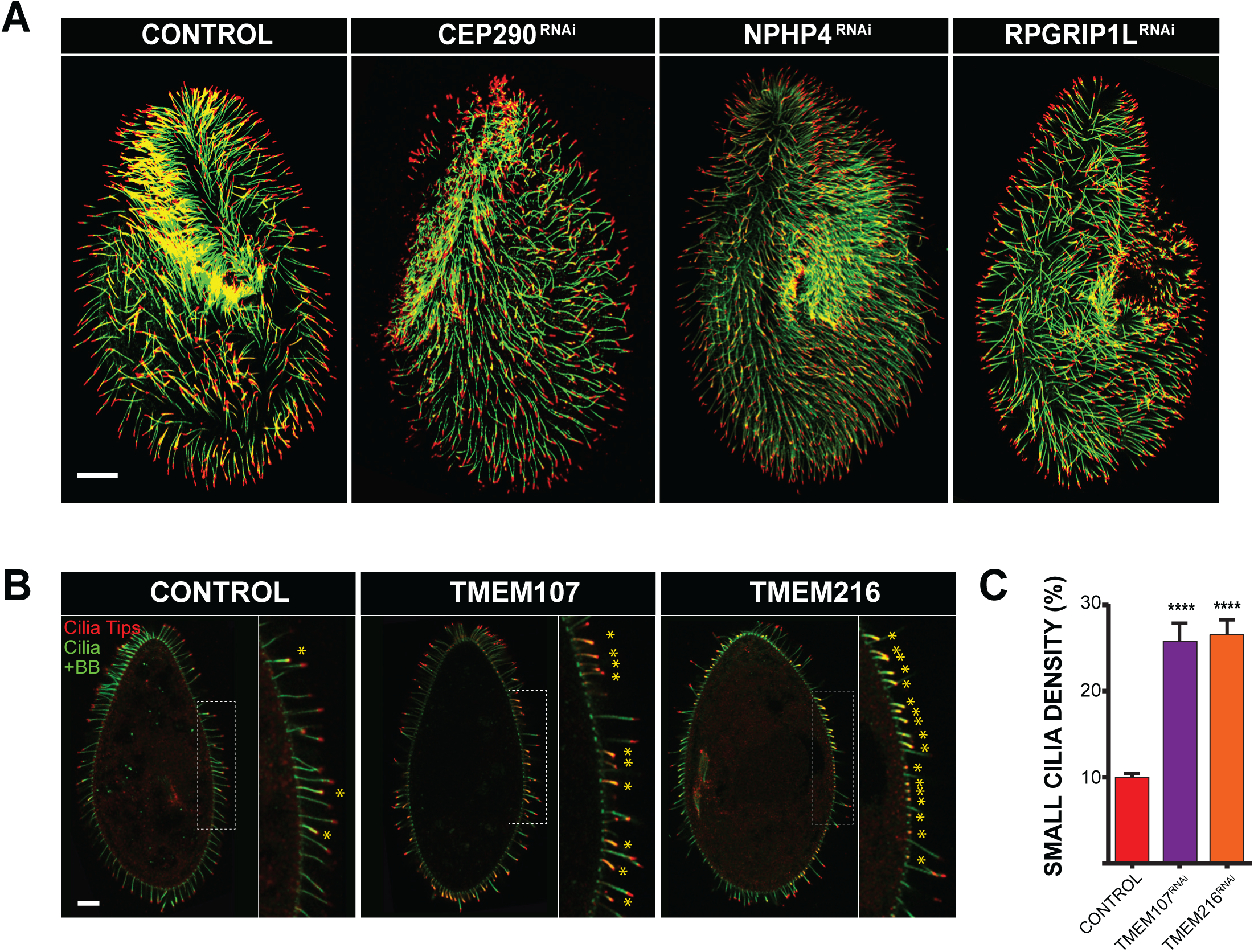
Ciliary pattern of paramecia depleted for TZ proteins. A- Ciliary pattern of paramecia treated with control RNAi or TZ^RNAi^ (CEP290, NPHP4 and RPGRIP1L). Cells were immuno-stained by the monoclonal anti-mono-glycylated tubulin, TAP952 (red, cilia tip labelling) and the polyclonal poly-glutamylated tubulin, polyE antibodies (green, decorating BB and cilia). These TZ depleted paramecia display a ciliary pattern similar to control paramecia. B- Control^RNAi^, TMEM107^RNAi^ and TMEM216^RNAi^ paramecia stained for cilia using TAP952 (red) and the polyclonal antibodies directed against poly-E tubulin (green). Control paramecia show the usual ciliary pattern with long cilia (10μm) and few short growing cilia indicated by an asterisk. A large increase in short (about 4mm) or tiny cilia is observed in TMEM107 and TMEM216 depleted cells. Bar =10μm **C-** Graph bars showing the mean percentage of small cilia in Control (n=60 cells), TMEM107 (n=50 cells) and TMEM216 (n=42 cells) depleted cells. Error bars show the SEM. Statistical significance was assessed by an unpaired t-test. ****(P<0.0001).

### TMEM107^RNAi^ and TMEM216^RNAi^ induce spontaneous deciliation

To first assess whether the ciliary phenotype observed in TMEM107^RNAi^ or TMEM216^RNAi^ cells could be caused by constant cilia breakage/regeneration, we searched for the presence of free detached cilia in the culture medium using IF staining (see Material and Methods). Although no cilia were found either in the medium of control cells or in CEP290^RNAi^, RPGRIP1L^RNAi^, NPHP4^RNAi^ cells, numerous cilia were detected in TMEM107^RNAi^ (mean number of 15.73 +/- 15.43 cilia per field) or TMEM216^RNAi^ (12.01+/-6.7) (Figure 3A). These results support the hypothesis that the depletion of TMEM107 or TMEM216 induces spontaneous cilia breakage, which was never seen in CEP290, NPHP4 or RPGRIP1L depleted cells or in controls.

**Figure 3:**
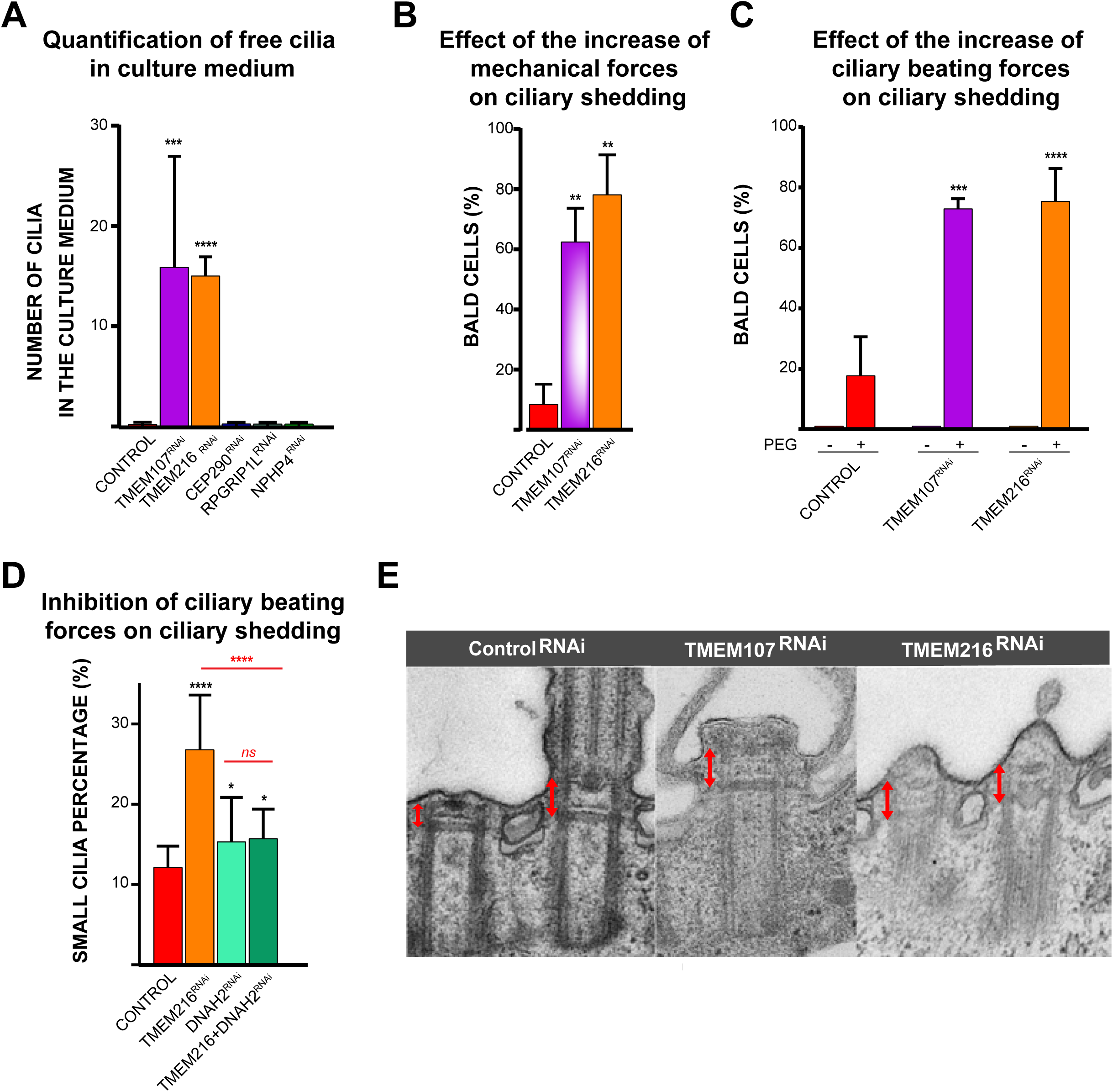
Ciliary shedding after TMEM107 or TMEM216 depletion. A- **Quantification of free cilia in culture medium**: Graph bars showing the mean number of free cilia found in the culture medium (n= 10 microscope fields analysed and 2 replicates). Error bars represent the standard deviation. Statistical significance was assessed by an unpaired t-test. ***(P=0.0004), ****(P<0.0001). B- **Effect of the increase of mechanical forces on ciliary shedding**. Left: Scheme depicting the experimental procedure, control Paramecia (labelled by India ink, in black) and TMEM107^RNAi^ or TMEM216^RNAi^ cells are aspirated (>10 up and down) in a glass micropipette in order to break fragile cilia. Cells were then fixed and stained for cilia. Right: quantification of the mean number of bald cells (<25% cilia per cell). N=3 replicates and > 100 cells per condition. Error bars represent the standard deviation. Statistical significance was assessed by an unpaired t-test. ** (0.0013<P<0.0021). C- **Effect of the increase of ciliary beating forces on ciliary shedding** Quantification of the mean number of bald cells (<25% cilia per cell) after 1H in PEG10% for control (n=410 cells), TMEM107^RNAi^ (n=157) and TMEM216^RNAI^ (n=303 cells). N=3 replicates. Errors bars represent the standard deviation. Statistical significance was assessed by an unpaired t-test. ***(P=0.0003) and **** (P<0.0001). D- **Inhibition of ciliary beating on ciliary shedding** Quantification of the mean cilia percentage in Control (n=60 cells), TMEM216^RNAi^ (n=42 cells), DNAH2^RNAI^ (n=15 cells) and TMEM216-DNAH2^RNAi^ (n=16 cells) *Paramecia*. DNAH2 cells present a small cilia percentage slightly higher than the controls. Impairing ciliary beating of TMEM216 cells by DNAH2 decreases the small cilia percentage. Statistical significance was assessed by unpaired t-test. * (P=0.0306), ** (P=0.0018) and **** (P<0.0001) and ns: non significant E- Representative EM images of ciliary defects induced by TMEM107^RNAI^ and TMEM216^RNAI^. Control^RNAi^ basal bodies showing a 2 basal body unit, with one unciliated and one ciliated BB. The length of the TZ is indicated by a red arrow. Cilia are either severed at the level of the axosomal plate as shown in TMEM107^RNAi^ or in a regrowth process as in TMEM216^RNAi^. Note that the length of the TZ corresponds to the length of TZ of ciliated BB, indicated by e red arrow.

To bring additional support to the hypothesis of constant breakage/regeneration we tested the resistance of cilia of RNAi-treated cells to a mechanical shearing. After ciliary immunolabelling, we observed that many TMEM107^RNAi^ and TMEM216^RNAi^ cells were bald, as defined by the loss of about 75 % of their cilia. The quantification of these bald cells in each RNAi condition confirmed that TMEM107^RNAi^ and TMEM216^RNAi^ cells deciliate easily: 62.4 +/-11.2 % of TMEM107^RNAi^ cells (n=116 cells in 3 replicates) and 78.1+/-13.3 %, TMEM216^RNAi^ cells (n=100) were bald versus 8.4+/-6.7 % for control cells (n=124; Figure 3B). Importantly, bald cells were never observed without this mechanical shearing. In summary, the presence of numerous tiny or short cilia stretches together with the occurrence of isolated cilia in the culture medium suggest that the depletion of either of the two TMEM proteins studied leads to spontaneous shedding of cilia. This result is confirmed by the ability of the TMEM depleted cells to deciliate easily after mechanical stress as compared to the control.

### Modification of the ciliary beating force influences ciliary shedding in TMEM107 and TMEM216 depleted cells

We hypothesized that the depletion of TMEM107 or TMEM216 weakened the cilia, which consequently did not resist the forces exerted to sustain ciliary beating or bending. One might expect in this situation that an increase of ciliary beating forces would exacerbate the deciliation phenotype while its decrease would alleviate it. To challenge this hypothesis, we first increased the paramecium standard-medium viscosity, known to induce an elevation of the ciliary force. For this, we transferred TMEM107^RNAi^ and TMEM216^RNAi^ cells or control paramecia to 10% PEG-containing standard culture medium for 1 hour. After fixation and IF, the ciliation pattern was analysed and bald cells were quantified (Figure 3C). This treatment affected control cells, since 16.92 +/- 12.74 % of them became bald (N=402 cells in 5 replicates). This percentage was nevertheless dramatically increased in TMEM107^RNAi^ (71.5+/-3.2%, N=157 cells in 2 replicates) and TMEM216^RNAi^ conditions (73.9+/-10.9%, N=303 cells in 5 replicates).

Conversely, we blocked the ciliary beating in TMEM216^RNAi^ cells by inactivating the gene encoding dynein axonemal heavy chain 2 (DNAH2), required for ciliary beating (Perrone *et al*, 2000). As expected, both the depletion of DNAH2 and co-depletion of DNAH2/TMEM216 led to a complete immobilization of the cells within 48h. As predicted, the co-depletion of TMEM216 and DNAH2 prevented ciliary breakage reducing it from 27% in TMEM216^RNAi^ to 15% (15.7+/- 0.9%, N=16 cells, see Figure 3D). Bald cells were never observed in these conditions. DNAH2 depletion alone induces a slight increase in the percentage of short cilia compared to control cells (15.31 +/- 1.4 %, N=15 cells versus 12%) (Figure 3D).

Altogether, these results demonstrate that TMEM107 and TMEM216 prevent cilia breaking against forces generated during ciliary beating.

### Cilia shedding occurs at the distal part of the transition zone, precisely at the axosomal plate level

To determine the potential site of cilia breakage in TMEM107^RNAi^ and TMEM216^RNAi^ cells, electron microscopy experiments were carried out. Control cells displayed the usual pattern of cortical two BB units showing a ciliated and an unciliated basal body, which differ in the size of the TZ, as previously reported (Tassin *et al*, 2015) (Supplemental Figure 1B). In cells depleted for TMEM107 or TMEM2016, numerous cilia showed a normal morphology, as expected from the IF staining. Nevertheless, BBs harbouring a typical deciliated structure (Adoutte *et al*, 1980) were detected: they displayed an extended TZ severed at its distal part just above the axosomal plate (Figure 3E TMEM107^RNAi^, Supplemental Figure 6). Interestingly, two BB units with two short growing cilia, characterised by the presence of electron dense material and extended TZ were also observed (Figure 3E, TMEM216^RNAi^, Supplemental Figure 6). Such a situation is never observed in control cells. Two BB units, either in the invariant field or in the mixed field (Figure S1), never grow their cilia simultaneously. Therefore, this feature suggests that at least one of them has been broken. Altogether, these results are in favour of a breaking point just above the distal end of the TZ, at the level of the axosomal plate, as in other cell model (Blum, 1971).

### TMEM216 depletion affects ciliary and membrane fusion gene expression

Cilia regeneration following the deciliation process is correlated with a modification of the ciliary gene expression profile (Arnaiz *et al*, 2009, 2010). We expected that the depletion of TMEM107 or TMEM216 should also modify ciliary gene expression. We thus undertook a transcriptomic analysis of TMEM216 depleted paramecia by RNA deep sequencing. We found that 4295 genes present a significant modification of their expression profile compared to controls (fold change of +/-2, suppl tables S2, supplemental Figure 7). The specific effect of silencing TMEM216 on the transcriptome was confirmed by comparison with the transcriptome of IFT57^RNAi^ cells (Shi *et al*, 2018) which shows only about 50 genes differentially expressed. No overlap was observed between TMEM216^RNAi^ and IFT57^RNAi^ (see supplemental Figure 7 A and B). Concerning TMEM216^RNAi^, one fourth of the misregulated genes (1079/4295) correspond to genes identified as potential ciliary genes by transcriptomics, proteomics and comparative genomic analyses (Arnaiz *et al*, 2009, Yano *et al*, 2013 and see Figure 4 and Tables S2,). This corresponds to an enrichment of ciliary genes compared to their representation in the complete genome (4585/39642). Remarkably, 535 genes identified in our RNAseq screen are also differentially expressed during the reciliation process, 95% of them (507) being up or down regulated in the same direction (426 genes are upregulated both in response to TMEM216 ^RNAi^ and during reciliation and 81 are down regulated). These results suggest that the depletion of TMEM216 mimics a reciliation process as expected from our previous results (cilia breakage and presence of short cilia), a phenotype completely different from the IFT57RNAi phenotype characterised by total disappearance of the cilia (Shi et al 2018).

**Figure 4:**
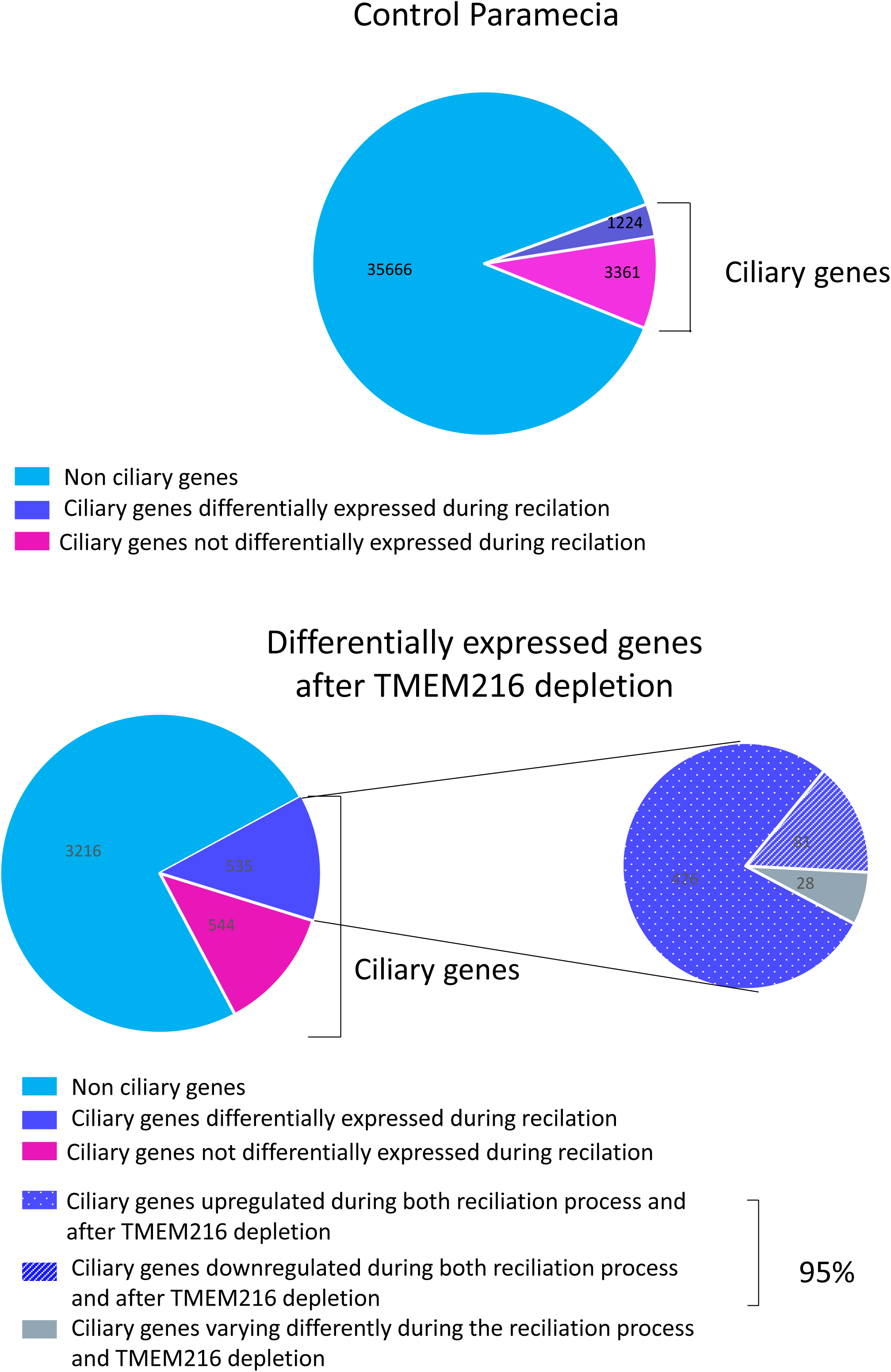
TMEM216/MKS2 RNAseq data. **A-** Diagram representing Paramecium genes in blue (40305 genes) of which roughly 10% are ciliary genes (4639). These ciliary genes were identified using the BioMart query tool of Cildb (Arnaiz et al. 2009). They correspond to genes whose expression is significantly modified during the reciliation process (stringency >=2 defined by Cildb) or encoding proteins identified in the ciliary or ciliary membrane proteome (stringency >=2) or whose homologs were recovered in at least 3 ciliary studies in other organisms. B- RNA seq analysis of paramecia depleted for TMEM216. Diagram represents genes differentially expressed after TMEM216^RNAi^. 4295 genes are differentially expressed and about ¼ of them are ciliary genes. 535 ciliary genes are differentially expressed during the reciliation process. Among these 535, 95% of them behave similarly (upregulation or downregulation) as during the reciliation process

We specifically searched for different proteins involved in several ciliogenesis processes such as transition zone formation, IFT transport, microtubule severing and membrane fusion. Except for the downregulation of the gene encoding the *Paramecium* B9D2 orthologs or IFT46 (fold change 0.48 Pvalue 3.54 E-4, fold change 0.21 Pvalue 3.8 E-3 respectively), the expression of proteins belonging to the MKS, NPHP, BBS or IFT complex identified in *Paramecium* was not affected. Concerning microtubule severing proteins, neither katanin nor spastin are differentially expressed during either chemical deciliation or TMEM216 depletion. In contrast, the expression of a VPS4 homolog and of a gene belonging to the SNF7 family, both involved in membrane fusion and shown to be associated with the TZ in *Chlamydomonas* (Diener *et al*, 2015, Ott *et al*, 2018), were significantly up-regulated (fold change 14.6, Pvalue 3.17 E-16 and fold change 21.8, P value 8 E-11 respectively). In addition, the expression of another gene involved in membrane fusion, NSF, is also greatly up regulated (fold change 40.59, P value 4.79 E-58). These proteins may act to remodel the cell membrane after axoneme severing.

### Depletion of CEP290, RPGRIP1L or NPHP4 affects the deciliation process

Although the depletion of CEP290, RPGRIP1L and NPHP4 did not modify the ciliary pattern, we wanted to assess whether it could affect the deciliation process induced by Ca^2+^/EtOH buffer. Indeed, CEP290^RNAi^, RPGRIP1L^RNAi^ or NPHP4^RNAi^ cells continued to swim after treatment while control cells immediately stopped moving, suggesting that deciliation was inefficient in these three RNAi conditions. Cilia immunostaining confirmed this assumption, since control cells appeared completely bald while CEP290-, RPGRIP1L- or NPHP4-depleted cells remained mostly ciliated (Figure 5A-B).

**Figure 5:**
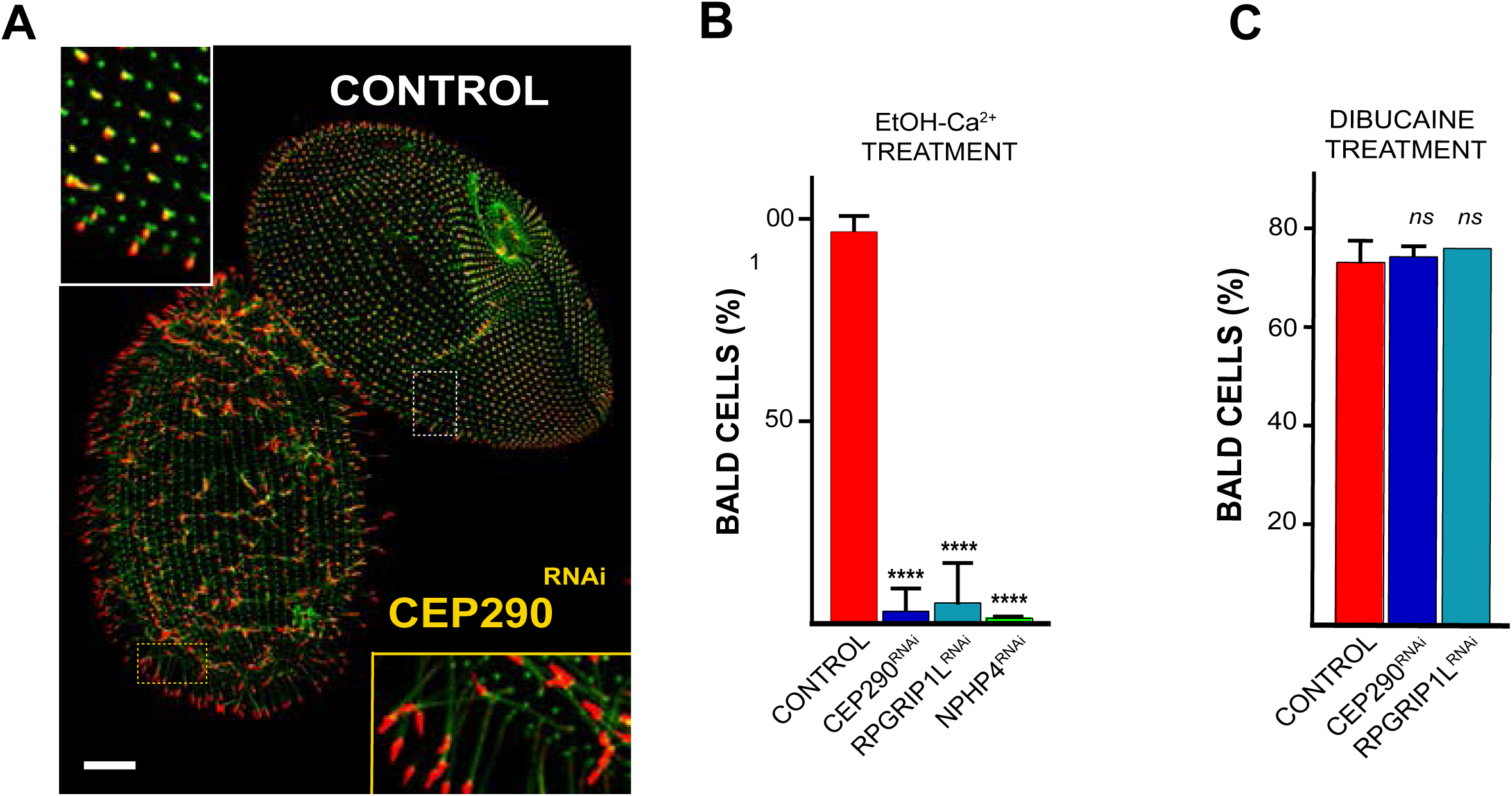
Depletion of CEP290, NPHP4 or RPGRIP1L alters deciliation ability. A- Control and CEP290 depleted cells were simultaneously submitted to deciliation treatment (5% EtOH, 1mM Ca^2+^) then labelled for cilia (TAP952 in red, polyE in green). Control cells were previously marked by India ink allowing their identification. Cep290^RNAi^ cells bear cilia whereas control cells are completely bald. Bar = 10 μm. B- Graph bars showing the mean percentages of ciliated cells (more than 50% of cilia) for control (n= 421 cells), CEP290^RNAi^ (n=98 cells), RPGRIP1L^RNAi^ (n=302 cells) and NPHP4^RNAi^ (n=82 cells) after deciliation by 5% EtOH, 1mM Ca^2+^. Error bars represent the standard deviation. 3 replicates per condition. Statistical significance was assessed by unpaired t-test. ****(P<0.0001). C- Graph bars showing the mean percentages of bald cells (less than 25% of cilia) for control (n=53 cells), CEP290^RNAi^ (n=47 cells) and RPGRIP1L^RNAi^ (n=68 cells) after deciliation with 25mM dibucaine. Error bars represent the standard deviation. 3 replicates per condition. Statistical significance was assessed by unpaired t-test. *ns*: non significant.

Studies at the molecular and genetic levels in *Chlamydomonas* showed that the deciliation process requires both a transient increase of intraciliary Ca^2+^ and functional microtubule severing proteins such as katanins and spastins (Lohret *et al*, 1999; Quarmby, 2009). To determine if the deciliation defects observed after depletion of CEP290 or RPGRIP1L were due to defective signal transduction or to inability of microtubule severing proteins to break the axoneme, we turned to a dibucaine treatment, which is known to increase intracellular and intraciliary calcium concentration by blocking Ca^2+^ ion extruding channels (Quarmby, 2004). When exposing RNAi-treated paramecia to 5mM dibucaine, they rapidly stopped swimming and became bald (Figure 5C). These results indicate that CEP290^RNAi^ or RPGRIP1L^RNAi^ cells possess functional microtubule-severing agents necessary for cilia shedding and that the defects observed are most likely due to a defective Calcium signalling pathway.

### The conserved domains (Coiled-coiled region and C2 domains) of RPGRIP1L are required for the deciliation process

The alignment of the *Paramecium* RPGRIP1L proteins with their orthologs showed that, in addition to the conserved N-terminal coiled-coiled region and central C2 domains (Zhang & Aravind, 2012), they display extra EF-hand domains instead of the RPGR binding domain present in the C-terminal part of the protein in all metazoans. The EF-hand domains are known to undergo conformational changes upon calcium binding. Since a defective Calcium signaling pathway could be the cause of the absence of cilia autotomy in RPGRIP1L-depleted cells, we wanted to determine if the effect of RPGRIP1L on the deciliation process might depend on these additional C-terminal EF hand domains or on the conserved N terminal region. For this, we introduced and expressed in paramecia a version of RPGRIP1L deprived of its EF-hand domains (RPGRIP1LΔEF), which localises like the full length protein (Supplemental Figure S8B), to see if it might rescue the RPGRIP1L depletion. Transformants expressing either GFP-tagged RPGRIP1LΔEF or full length RPGRIP1L were treated with RNAi sequences targeting EF-hand domains (depleting only endogenous and full length RPGRIPL-GFP) (see Schema Figure S8C). If the conserved N-terminal domains are required for the deciliation process, a complementation should be observed only on transformants expressing RPGRIP1LΔEF when EF-hand RNAi targeting sequence is used, and cells should deciliate like controls. Our results (supplemental Figure 8C, D) confirm this hypothesis and suggest that this function could therefore be conserved in metazoans.

### Depletion of RPGRIP1L, NPHP4 or CEP290 affects TZ structure

In order to determine whether ultrastructural defects are associated with RPGRIP1L, NPHP4 or CEP290 depletion, we undertook electron microscopy analyses. Even if most of the cilia appeared normal, some atypical cilia were observed in NPHP4 depleted cells (Figure 6B). This is reminiscent of the weak ultrastructural defects observed in NPHP4 mutants of other cell model organisms (Williams et al 2011, Awata et al 2014). RPGRIP1L depleted cells also displayed aberrant cilia showing developed TZ with clear lack of connections between the TZ region and the membrane (Figure 6C, D). Cilia displaying an amazingly enlarged lumen in CEP290 depleted cells (Figure 6E, F), were also observed as in *Chlamydomonas* (Craige *et al*, 2010) and in multiciliated cells of non-syndromic Leber congenital amaurosis patients patients harbouring *Cep290* mutations (Papon *et al*, 2010). In agreement with IF data, BB anchoring was achieved normally. Altogether, these data suggest that these electron microscopy phenotypes are similar to the ones observed in other cellular models suggesting that the function of these proteins is conserved in *Paramecium*.

**Figure 6:**
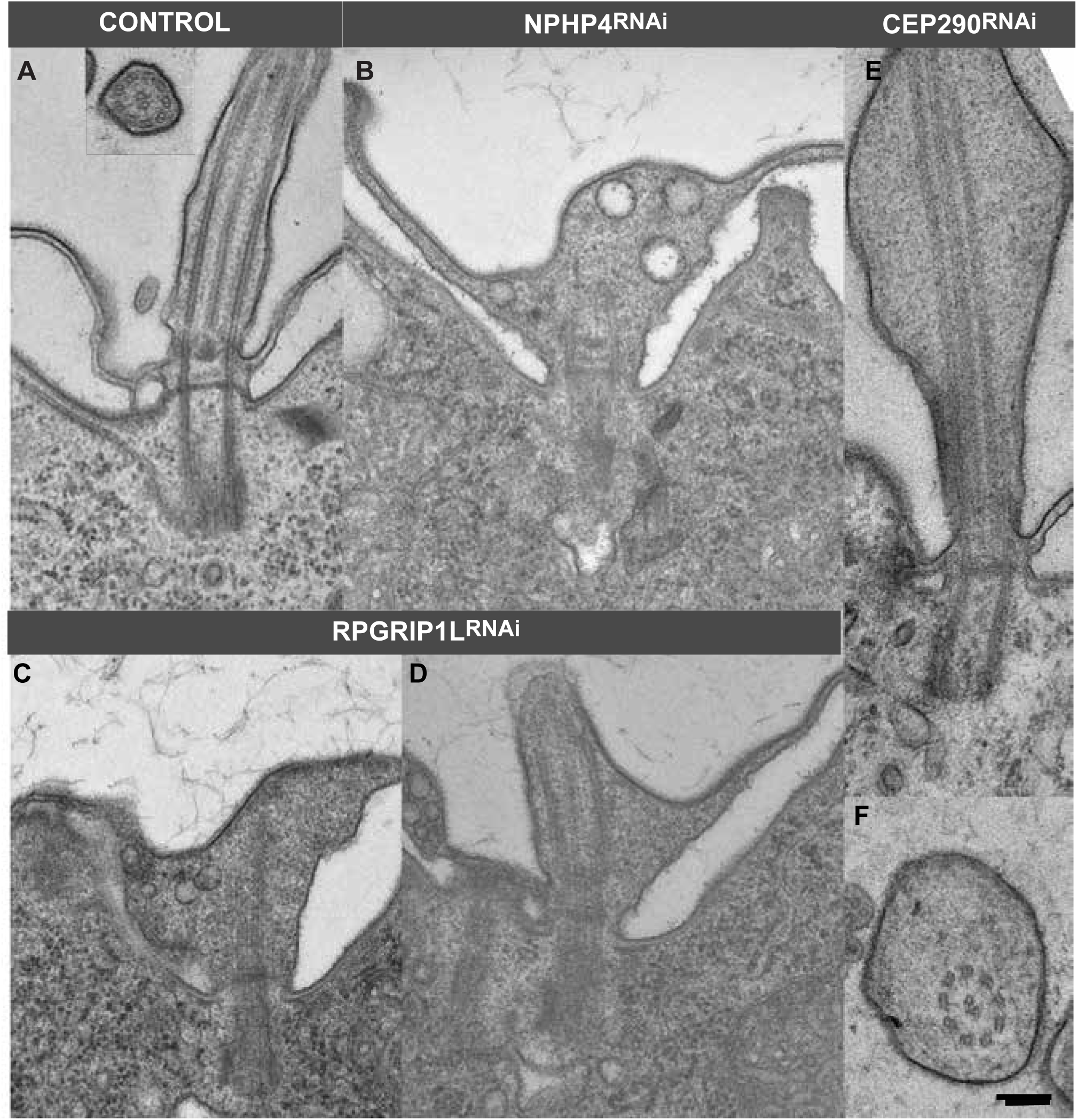
Depletion of CEP290, NPHP4 and RPGRIP1L affects ciliary shape. Electron microscopy images of abnormal cilia found in NPHP4^RNAi^ (A), RPGRIP1L^RNAi^ (C-D) and CEP290^RNAi^ (F-G), A control cilium is shown longitudinally as well as transversely in A. In NPHP4 and RPGRIP1L depleted cells, the link between the axoneme and the ciliary membrane seems destroyed at the level of the transition zone. Abnormal cilia present extension defects and vesicles accumulate inside the cilia, probably resulting from defective TZ gate function. In CEP290 depleted cells, abnormal cilia presenting a strikingly enlarged lumen, resulting most probably from altered ciliary gate function. Bar: 200nm.

## DISCUSSION

By studying their localisation as well as the effect of their depletion, we demonstrate that, in addition to their documented role in maintaining the ciliary gate structure and function, some proteins belonging to the MKS and NPHP complexes, as well as the scaffolding proteins RPGRIP1L and CEP290, play crucial and opposite functions in ciliary shedding in *Paramecium*.

As in other species (Yang *et al*, 2015, Jana *et al*, 2018, Lambacher *et al*, 2016), the studied *Paramecium* TZ proteins localise with nine-fold symmetry in several concentric rings located between the tubulin scaffold and the ciliary. All of them localise along the same axial axis at the level of the axosomal plate, a region closely associated to the ciliary necklace (Dute & Kung, 1978). Such localisation is in agreement with the assumption that CEP290, in association with MKS and NPHP modules, organizes the Y-links and the ciliary necklace at the TZ (Craige et al 2010, Williams et al 2011). This is reminiscent of the localisation on the same axial plane of RPGRIP1L and members of the MKS complex observed in RPE1 primary cilia, while in this case CEP290 is found underneath (Yang *et al*, 2015). However, it contrasts with the observations performed on cilia *in Chlamydomonas* (Craige *et al*, 2010, Awata *et al*, 2014), *Drosophila* (Jana *et al*, 2018) and *C. elegans* (Lambacher *et al*, 2016), in which CEP290, TMEM107 and TMEM216 extend along the longitudinal length of the TZ. These differences already noticed between species (Gonçalves & Pelletier, 2017) might reflect the differences in organization of the TZ, which could vary even from one cell type to another in the same organism (Wiegering *et al*, 2018a). Only two rows constitute the ciliary necklace in *Paramecium* explaining the localisation of the proteins at the same axial level (Dute and Kung 1980).

Depletion of either RPGRIP1L, NPHP4 or CEP290 proteins in *Paramecium* could induce defects in the structural integrity of the TZ as in other cell models. In the case of RPGRIP1L or NPHP4 gene silencing, clear disconnections between the TZ region and the membrane are observed indicating that they could have a role in tethering the axonemal microtubules to the ciliary membrane. As observed previously in nasal cilia of some non-syndromic Leber congenital amaurosis patients (Papon *et al*, 2010), CEP290 depleted cells display enlarged cilia, consistent with aberrant gating at the TZ.

By contrast the depletion of the two MKS complex proteins TMEM216 and TMEM107 does not seem to severely affect the ciliary and the TZ structures. This result is similar to the one observed in other systems such as *C elegans,* in which loss of function of *Tmem107* or *Tmem216* does not lead to gross structural or functional defects as judged by normal dye-filling (Lambacher *et al*, 2016, Huang *et al*, 2011).

In addition, during this study we discovered a novel important function of these proteins in the deciliation process. This was established by the fact that TMEM216 and TMEM107 depleted cells spontaneously shed their cilia into the extracellular medium by autotomy, followed by a ciliary regrowth, whereas RPGRIP1L, CEP290 and NPHP4 by contrast are resistant to deciliation induced by Ca2+/ethanol buffer. Although we cannot totally exclude a contribution of ciliary involution, our data are totally consistent with a process of spontaneous deciliation: i) numerous stretches of tiny or short cilia in the process of growing were observed on the surface of depleted cells (see Figure 2B); ii) detached cilia were recovered in the culture medium in which depleted cells were grown (graph Figure 3A);.iii) transcriptomic analysis reveals that ¼ of the genes differentially regulated after TMEM216 depletion are ciliary genes, half of them also being differentially regulated during the reciliation process, and 95% of them behave in the same manner (see Figure 4). Together with the results of modulation of ciliary beating experiments (Figure C, D), these data lead us to propose that the depletion of either TMEM107 or TMEM216 induces a weakening of the distal part of the TZ, which does not resist the shear stress induced by ciliary movements. Interestingly, this finding is consistent with the TZ weakening observed in primary cilia depleted for TCTN2, another transition zone protein (Weng *et al*, 2018), necessary for the localisation of both TMEM216 and TMEM107.

As described above the cilia of RPGRIP1L, NPHP4 or CEP290 depleted cells are resistant to Ca2+/ EtOH-induced deciliation but shed their cilia during dibucaine treatment indicating that microtubule severing is not affected in these cells.

Little is known about the deflagellation process that requires both outer microtubule-severing and membrane pinch off at the distal part of the TZ. However, the work of Quarmby and collaborators in *Chlamydomonas* (Quarmby, 2004, Quarmby, 2009) shed some light on this process, which requires a Ca^2+^ influx. Several screens using acidic stress have allowed selection of deflagellation-defective mutants which can be separated in two groups (Finst *et al*, 1998, Hilton *et al*, 2016). The first one is constituted of mutants supposed to be defective in the microtubule-severing events. It includes mutants in the *Fa1* and *Fa2* genes which encode respectively a Ca^2+/^calmodulin containing protein (Finst *et al*, 2000) and a NIMA related kinase (Mahjoub *et al*, 2002). The second class of mutants (*ADF*) is constituted of strains which are supposed to be affected in the pathway that activates a calcium influx involved in deflagellation (Quarmby & Hartzell, 1994, Quarmby, 1996) : *ADF1* encodes a TRP Ca^2+^ channel and *ADF3* a conserved microtubule binding protein FAP16, (Hilton *et al*, 2016). Interestingly, these proteins localised either to the TZ or just above at the very proximal part of the cilia. Since the depletion of CEP290, RPGRIP1L or NPHP4 disturbs TZ integrity in *Paramecium*, this defect might be accompanied by a mislocalisation of proteins involved in the deciliation process such as the TRP calcium channel or FAP16 as described in C*hlamydomonas*.

The regulation of phosphoinositide levels at the TZ has also been involved in flagellar. Indeed in *Chlamydomonas*, it has been shown that phosphatidylinositol 4,5 biphosphate PI(4,5)2 (PIP2) and its hydrolysis products were involved in the deflagellation process. First, the ADF2 gene could encode an inositol 1,3,4-triphosphate 5,6 kinase although this is not fully demonstrated (Hilton *et al*, 2016). Second, Ca^2+^ entry during the deflagellation process induces the activation of phospholipase C that triggers the hydrolysis of PIP2 into inositol 1,4,5 triphosphate I(1,4,5)P3 and DAG (Quarmby *et al*, 1992, Yueh & Crain, 1993, Quarmby & Hartzell, 1994).

Recent studies in mammalian cells points to the accumulation of PIP2 in cilia as a « cut here signal » which leads to cilia decapitation upon mitotic entry (Phua *et al*, 2017). Enriched at the ciliary base in resting cells, the concentration of PIP2 progressively decreases along the TZ and is excluded from the cilia due to its conversion to PIP by INPP5E, an inositol polyphosphate 5-phosphatase localised in the cilia (Chávez *et al*, 2015, Garcia-Gonzalo *et al*, 2015). Upon mitotic entry, PIP2 accumulates at the distal end of cilia, inducing actin polymerization, which is required for membrane scission. The position of this cilia excision signal is correlated with the maximal ciliary PIP2 accumulation. Therefore, as already suggested by Mirvis *et al*, (2019) who saw primary cilia shedding during primary cilium disassembly, we propose that a modification of the PIP2 concentration at the level of the TZ could trigger the ciliary autotomy process, PIP_2_ distribution being controlled by INPP5e, whose enrichment relies on the integrity of the TZ. Inositol lipids are found in *Paramecium* ciliary membranes together with the kinase and phosphatase that interconvert PI, PIP and PIP2 (Suchard *et al*, 1989). One can thus suppose that TMEM216 and TMEM107 depletion could affect the distribution of INPP5E, which in turn leads to accumulation of PIP2 and membrane scission. A differential modification of the PIP2 content along the cilia in CEP290, NPHP4 or RPGRIP1L-depleted cells compared to the TMEM216 or TMEM107-depleted ones might explain the difference in the resistance to deciliation observed between the different cells.

Despite being particularly well described in unicellular organism, physiological deciliation appears to be a more widespread process than previously thought. In metazoans, deciliation has been observed in various mammalian species during the menstrual cycle as for example the monkey oviduct (Brenner, 1969) and following castration in rabbit (Rumery & Eddy, 1974). Boisvieux-Ulrich (1980), studying the deciliation in quail oviduct after progesterone treatment, described two types of mechanisms responsible of the process. The first type is a resorption of the entire axoneme into the cell; the second is the shedding of cilia into the magnum lumen, which arises by an alteration of the TZ neck region. The latter mechanism occurs on individual or fused cilia, in which the TZ remains individual while the ciliary membrane fuses along the entire length, producing a polyaxoneme within a single membrane. Deciliation in metazoans is not a specific fate of multiciliated cells. Epithelial cells could be reversibly deciliated by diverse agents (Praetorius & Spring, 2003, Overgaard *et al*, 2009). Recently, an analysis of primary cilium disassembly by long-term live cell imaging indicated that its occurs predominantly by cilia shedding (Mirvis *et al*, 2019). Finally, apical abscission separating the centrosome from the cilium has been observed during neurogenesis, allowing the detachment of the neurons from the ventricle in order to migrate to the lateral neural tube (Das & Storey, 2014).

All these results demonstrate that cilia autotomy is a general process occurring in numerous eukaryotes either physiologically or under stress or pharmacological conditions. Our results lead us to propose that in mammals the regulation of distal TZ molecular content by MKS/NPHP genes is crucial for the cells to trigger cilia autotomy with an appropriate timing.

## MATERIAL AND METHODS

### Strains and culture conditions

Stock d4-2 of *P. tetraurelia*, the wild-type reference strain, was used in all feeding experiments. The nd7-1 mutant, carrying a recessive monogenic mutation preventing trichocyst discharge (Skouri & Cohen, 1997), was used for the expression of GFP-fusions. Cells were grown at 27°C in a wheatgrass infusion (BHB, GSE Vertrieb, Germany), bacterized with *Klebsiella pneumoniae*, and supplemented with 0.8 μg/ml β-sitosterol according to standard procedures ((Sonneborn, 1970)).

### Gene identification

Genes were identified using CilDB (http://cildb.i2bc.paris-saclay.fr/, Arnaiz et al, 2009, 2014) and by BLAST search in ParameciumDB (https://paramecium.i2bc.paris-saclay.fr/ Arnaiz et al, 2011). The complete list of genes identified is given in TableS1. Tophat2 (v2.0.12 –min-intron-length 15 –max-intron-length 100) was used to map paired-end reads (2 replicates) on the *P tetraurelia* MAC reference (ptetraurelia_mac_51.fa). Differential gene expression analysis was done with DESeq2 (v1.4.1) package. Raw reads were deposited in the ENA under the project accession PRJXXX.

### Gene cloning

Genomic DNA was amplified by PCR, using specific primers (given in table S3) in order to fuse the coding regions of TZ genes to the GFP coding sequence in the pPXV vector, between *Spe*I and *Xho*I restriction sites. If these restriction sites were present in the sequence to be cloned, we used the Gibson cloning method (Gibson *et al*, 2009). For gene silencing constructs, sequences of interest, with the help of the RNAi off target tools in ParameciumDB (Arnaiz et al, 2007), for their ability to inactivate all the ohnologs, when possible, and to prevent RNAi off target. The RNAi sequences were cloned into the feeding vector, L4440, between two T7 promoters (Timmons & Fire, 1998).

### Paramecium transformation

*Nd7-1* cells were transformed by microinjection into their macronucleus (Gilley *et al*, 1988). DNA containing a mixture of the plasmid of interest (5 μg/μl) and of plasmid DNA directing the expression of the *ND7* gene. Microinjection was made under an inverted Nikon phase-contrast microscope, using a Narishige micromanipulation device and an Eppendorf air pressure microinjector. Transformants were screened for their ability to discharge their trichocysts and further analyzed for GFP.

### Gene silencing

L4440 constructs were used to transform HT115 bacteria, allowing the production of double-stranded RNA. WT cells were fed with these bacteria and transferred daily into fresh medium at 27°C (TZ*^RNAi^*) (Galvani & Sperling, 2002). Control cells were fed with HT115 bacteria carrying the L4440 vector containing the *ND7* gene.

### Fluorescence microscopy

In order to ascertain that our immunostaining protocol was not affecting ciliation pattern, ink-labeled controls were always mixed with RNAi-treated cells and all cells treated simultaneously within the same dish. India ink was added to control cultures at least 30 minutes before staining. India ink particles are nontoxic for the cells, but are absorbed in the feeding vacuoles, which allows detection of control cells by their dark vacuoles using phase contrast. The unlabeled cells are referred to TZ^RNAi^.

Cilia are often more fragile when paramecia are coming from 27°C, so cells had to be kept 1 or 2 h at room T° before proceeding to IF.

50-100 cells were collected in the smallest volume possible and were permeabilised in 200μl PHEM (Pipes 60mM, Hepes 25 mM, EGTA 10mM, MgCl2 2mM, adjusted to pH 6.9) with 0.5% saponin for 30 seconds. An equal volume of PHEM, 2% PFA, 0.5% saponin was added for 15mn. Buffer was then aspirated and cells were rinsed 3 times for 10 minutes in PHEM, 0.5% saponin. Monoclonal antibodies diluted in PHEM, 0.5% saponin were incubated for 15 mn. After 3 washes, cells were incubated with the appropriate secondary antibodies from Thermofisher at a dilution of 1:500 together with DAPI. A final wash was done and cells were mounted in citifluor AF1 (EMS, Hatfield, PA). Confocal acquisitions were made with a Leica SP8 equipped with a UV diode (line 405), and three laser diodes (lines 488, 552 and 635) for excitation and two PMT detectors. Image stacks were processed with imageJ and Photoshop. See (Aubusson-Fleury *et al*, 2015) for IF methods, antibodies specificity and cilia staining. The following antibodies were used: The monoclonal anti–tubulin ID5 (1:100) recognizes poly-glutamylation of α-tubulin (Wehland & Weber, 1987, Rüdiger *et al*, 1999) a gift from J. Wehland; the monoclonal TAP952 antibodies (1:50) recognizes monoglycylation (Callen *et al*, 1994), the polyclonal PolyE (1:100) anti-tubulin (Janke & Bulinski, 2011); the polyclonal anti-GFP antibodies (1:1000, Interchim, Montlucon, France).

### STED analysis

STED imaging was performed using a Leica TCS SP8 STED 3X (Leica Microsystems CMS GmbH, Mannheim, Germany). The system was equipped with a WLL ranging from 470 to 670 nm for excitation and with 3D STED lasers at 592, 660, and 775 nm. A 100× 1.4 Oil STED white objective was used to acquire the images. GFP, AF568, and Cy5 were excited at 488, 561, and 643 nm, respectively. Detection ranges were 500–550, 575–625, and 660–700 nm, respectively. A pixel size of 25 nm was used. For deconvolution, SVI Huygens was used. Diameters were measured using the plot profile tool in imageJ to obtain the distance between the 2 maximum intensity peaks.

### Electron microscopy

For ultrastructural observations, cells were fixed in 1% (v/v) glutaraldehyde and 1% OsO4 (v/v) in 0.05 M cacodylate buffer, pH 7.4 for 30 min. After rinsing, cells were embedded in 2% agarose. Agarose blocks were then dehydrated in graded series of ethanol and propylene oxide and embedded in Epon812 (TAAB, Aldermaston, Bershire, UK). For pre-embedding immunolocalisation, the immunostaining process was carried out as described for immunofluorescence using gold-coupled instead of fluorochrome-coupled secondary antibodies (gold-labeled anti-rabbit IgG -GAR G10, Aurion) diluted 1/50 for 30 mn. Cells were then treated as described above.

All ultrathin sections were contrasted with uranyl acetate and lead citrate. The sections were examined with a Jeol transmission electron microscope1400 (at 120kV)

### Swimming analysis

4 to 8 Paramecia were transferred in 10 μl drops of conditioned BHB (bacterized BHB solution, then depleted of bacteria after their growth and sterilized) for 15 minutes before being tracked for 10s every 0.3s. We used a Zeiss Steni 2000-C dissecting microscope with 1-min time-lapse acquisitions at 7 frames per second with a Roper Coolsnap-CF camera and Metamorph software (Universal Imaging). Stacks were analyzed using the Manual tracking tool in ImageJ.

### Quantification of free cilia in culture medium

After 24h of gene silencing, cells were transferred into fresh medium in order to obtain 8 cells per 300 μl well at t=48h. The number of cells was adjusted according to the dividing capacity of the RNAi condition. At 48h of RNAi treatment, all the cells were carefully aspirated from the culture wells in order to collect the amount of medium corresponding to 300 cells (approximately 30-35 wells per condition). The culture medium was then centrifuged onto a microscopy coverslip as previously described for centrosomes (Gogendeau *et al*, 2015). Cilia were fixed with Methanol-20°C for 6 minutes and stained using TAP952 and polyE antibodies. The number of cilia per microscope field (184.7×184.7μm) was quantified.

### Deciliation ability analysis

We measured cilia resistance to deciliation in two different ways. First, cells, RNAi conditions together with ink-labelled controls, were submitted to vigorous pipetting using a thin glass micropipette (>10 up and down movements). In the second test, cells were placed for 1 hour in 10% PEG6000 (Merck) (diluted in conditioned BHB). Cells were then directly submitted to immunolabelling and observed under the microscope. Cells were classified into 4 categories according to their ciliation state: Normal, 75% cilia (= few holes in the ciliation), 50% cilia and bald (less than 25% of cilia).

### Deciliation experiments

Standard protocol: paramecia (RNAi-treated and ink-labelled controls) were transferred to 1ml deciliation buffer (10 mM Tris, 1 mM Ca^2+^, 5% EtOH) and vigorously vortexed for 2 minutes to allow the cilia to shed.

Dibucaine treatment: cells were resuspended in a solution containing 5 mM MgSO4, 4% sucrose and 25 mM Hepes. Dibucaine-HCl (Sigma) 25m M was added to obtain a final concentration of 5 mM according to Nelson (Nelson, 1995)

After deciliation treatment, cells were rinsed and immuno-stained for cilia.

---

**Figure S1: Ciliation status in *Paramecium***

A- Paramecia labelled by the monoclonal anti-glutamylated tubulin ID5 (decorating basal bodies (BB)). Paramecium cortex presents different regions in which the basal body pattern differs. The red star highlights the invariant field region where BB are organised in doublets. In this field, each BB of the doublets is ciliated. The mixed field, highlighted here in light grey, presents an alternation of BB singlets and doublets. In the mixed field, only the posterior BB bears a cilium and the anterior one remains unciliated until the next cell cycle. The region in white only possesses singlet BBs. The singlet BBs are not all ciliated at the exit of mitosis. However, *Paramecium* cells will grow cilia all along the cell cycle and all singlet BBs will bear a cilium before the next mitosis. Bars= 10μm and 1μm respectively.

B- Electron microscopy images showing a ciliated doublet in the invariant field (left) and a doublet in the mixed field, where only one BB is ciliated and the other one is anchored but not ciliated (right). The transition zone is characterised by the presence of 3 successive layers indicated by black arrowheads, from bottom to top: the terminal plate, the intermediate plate and the axonemal plate. Strikingly, transition zone evolves structurally between non-ciliated and ciliated BB. The length of the TZ is indicated by red arrows. Note that the unciliated BB in the mixed field shows a reduced TZ compared to ciliated ones. Bar=200nm.

**Figure S2: *Paramecium* Transition zone proteins are recruited when cilia extend**

A- Localisation of TZ-GFP fusions when Paramecia were fixed before permeabilization. Labelling using a mix of the monoclonal anti-glutamylated tubulin ID5 (decorating basal bodies (BB) and cilia in red), the monoclonal anti-polyglycylated tubulin, Axo49 (decorating cilia, also in red) and a polyclonal anti-GFP (in green). TMEM216-GFP localises both at the transition zone of ciliated basal bodies and at the proximal side of non-ciliated ones. This proximal signal completely disappears in cells permeabilised before fixation (See Figure 1A). Bar=1 μm.

B- Images of cilia at different steps of growth labelled by ID5, and anti-GFP. Cilia at different steps of their growth labelled by ID5 (which decorated also the basal body). To ascertain that cilia were growing and not broken, dividing cells where new cilia are generated were observed. To avoid fluctuations in fluorescence intensity due to variations in expression rate, all the cilia shown were extracted from the same cell for each GFP-tagged expression. The GFP signal is detected at the TZ as soon as the growing cilium is detected.

**Figure S3: Efficiency of inactivation of the different RNAi vectors**

The efficiency of the inactivation of the different genes is evaluated by decrease of the GFP fluorescence after 24 h of RNAi. For each TZ GFP transformants, a cell were fed with bacteria expressing control vector (left panel) and a cell fed with bacteria expressing TZ^RNAi^ (right panel) are shown. Note the decrease of the GFP labelling in each condition.

**Figure S4: Swimming behaviour of control and TZ^RNAi^**

Dot plot graph depicting the mean swimming speeds of control *Paramecia* and cells depleted by TZ genes for 24h and 48h. Each dot shows the mean velocity of 1 cell (n= 3 replicates and >50 cells per conditions). The lines represent the mean and the error bars the standard deviation. Statistical significance was assessed by unpaired t-test. ****(P<0.0001).

**Figure S5: Depletion of TZ proteins do not affect Basal body positioning**

Paramecia were decorated for basal bodies and cilia by the the polyclonal poly-glutamylated tubulin. Basal bodies are perfectly aligned along ciliary rows indicating an absence of BB duplication or anchoring defects. Bar=15μm

**Figure S6: TMEM107 and TMEM216 depleted cells shed their cilia distally of the transition zone**

Other example of shed cilia after the depletion of either TMEM107 or TMEM216. The transition zone indicated by a red arrow shows the length of TZ in agreement with a ciliated basal body. This indicates that the cilia have been shed.

**Figure S7: Heatmaps of differentially expressed genes identified in transcriptomic analyses of TMEM216^RNAi^ cells compared to control^RNAi^ and IFT57^RNAi^ cells.**

Less expressed genes are in blue and highly ones in red after TMEM216 expression. The top panel shows down-regulated genes and the bottom the up-regulated genes. Note that the differentially expressed genes identified in TMEM216 depleted cells are specific.

**Figure S8: RPGRIP1L EF-Hand domains are not involved in the deciliation signal.**

A- Paramecia expressing RPGRIP1L full length (FL, left) and RPGRIP1L short form (SF). These two constructs present exactly the same localisation. Bar= 10 μm.

B- Experimental design: WT paramecia were transfected either by RPGRIP1L-GFP full length or by RPGRIP1L-GFP short form deprived of EF-hand domains. Cells were then subjected to RNAi targeting only full-length forms (EF-hand RNAi target)). In cells expressing RPGRIP1L-GFP full length, should resist deciliation. The red cross on the protein schemas indicate that the protein will not be produced due to the RNAi. In cells expressing RPGRIP1LΔEF-GFP, EF-hand RNAi will only target the endogenous form

C- Quantification of deciliation experiment in RPGRIP1L-FL expressing cells (left) or RPGRIP1LΔEF expressing cells (right).

Left: Graph bars showing the mean percentages of ciliated cells (more than 50% of cilia) for control (n= 154 cells), EF-Hand^RNAi^ (n=96 cells). Error bars represent the standard deviation. 2 replicates per condition. Statistical significance was assessed by unpaired t-test. * (0.0121<P<0.0211).

Right: Graph bars showing the mean percentages of ciliated cells (more than 50% of cilia) for control (n= 62 cells), EF-Hand^RNAi^ (n=97 cells)). Errors bars represent the standard deviation. 2 replicates per condition. Statistical significance was assessed by unpaired t-test. *** (P=0.0003), *ns*= non significant.

**Table S1: ID of transition zone proteins identified in *Paramecium***

**Table S2: List of genes differentially expressed in silenced TMEM216 cells**

**Sheet 1**: *Paramecium* genes whose expression was significantly modified in TMEM216^RNAi^ cells. The genes mentioned in the text are in yellow.

**Sheet 2:** *Paramecium* genes whose the expression is significantly modified both in TMEM216^RNAi^ cells and during reciliation.

**Sheet 3:** Potential ciliary genes identified by proteomic and comparative genomic analyzes whose expression is significantly modified in TMEM216^RNAi^ cells

**Sheet 4:** *Paramecium* genes, not identified as ciliary genes, whose expression is significantly modified in TMEM216^RNAi^ cells

ID: ParameciumDB accession number

FC_TMEM216: Fold change value compared to the control.

NB_ciliary_evidence: number of high throughput ciliary studies in which the gene or its homologs were identified as a potential ciliary gene.

NB_PEP_Yano: number of peptides assigned to the encoded *protein* identified in the proteome of the ciliary membrane in *Paramecium* (Yano et al., 2013). NA: no peptide identified

Ciliary proteome: peptides assigned to the protein encoded by the gene were identified in the *Paramecium* ciliary proteome (Arnaiz et al.,2009)

Reciliation_transcriptome_significant: expression of the gene was significantly modified during the recilation process (Arnaiz et al., 2010)

Reciliation_transcriptome_cluster_att. The genes differentially expressed during reciliation were clustered according to their expression profile (Arnaiz et al 2010). This column indicates the cluster to which the gene was assigned. No cluster indicates that its expression was not significantly modified.

ANNOTATION: predicted function of the protein

interpro_desc_terms: InterPro predicted function of the protein

**Table S3: oligos used for cloning**

**Supplemental movies 1 to 6: trajectories of Paramecia under different RNAi conditions.**

Cells were imaged every 0.3 seconds during 10 seconds.

## ACKNOWLEDGEMENTS

We acknowledge the High-throughput sequencing facility of I2BC for its sequencing and bioinformatics expertise. The present work has benefited from Imagerie-Gif core facility supported by l’Agence Nationale de la Recherche (ANR-11-EQPX-0029/Morphoscope, ANR-10-INBS-04/FranceBioImaging; ANR-11-IDEX-0003-02/ Saclay Plant Sciences). We thank Cindy Mathon and Pascaline Tirand for excellent technical assistance. We tank Linda Sperling, Mireille Bétermier and all members of the Bétermier lab for stimulating discussions, Paul Guichard, Virginie Hamel and Thibaut Eguether for the critical reading of the manuscript and their suggestions.

The project has been funded by ANR *ANR*-10-*BLAN*-*1122 FOETOCILPATH* to J C and the ANR ANR-15-CE11-0002-01 (ANCHOR) to AMT.

## BIBLIOGRAPHY

Adoutte A, Ramanathan R, Lewis RM, Dute RR, Ling KY, Kung C & Nelson DL (1980) Biochemical studies of the excitable membrane of Paramecium tetraurelia. III. Proteins of cilia and ciliary membranes. J. Cell Biol. 84: 717–738

Arnaiz O, Goût J-F, Bétermier M, Bouhouche K, Cohen J, Duret L, Kapusta A, Meyer E & Sperling L (2010) Gene expression in a paleopolyploid: a transcriptome resource for the ciliate Paramecium tetraurelia. BMC Genomics 11: 547

Arnaiz O, Malinowska A, Klotz C, Sperling L, Dadlez M, Koll F & Cohen J (2009) Cildb: a knowledgebase for centrosomes and cilia. Database J. Biol. Databases Curation 2009: bap022

Aubusson-Fleury A, Cohen J & Lemullois M (2015) Ciliary heterogeneity within a single cell: the Paramecium model. Methods Cell Biol. 127: 457–485

Aubusson-Fleury A, Lemullois M, de Loubresse NG, Laligné C, Cohen J, Rosnet O, Jerka-Dziadosz M, Beisson J & Koll F (2012) The conserved centrosomal protein FOR20 is required for assembly of the transition zone and basal body docking at the cell surface. J. Cell Sci. 125: 4395–4404

Aury J-M, Jaillon O, Duret L, Noel B, Jubin C, Porcel BM, Ségurens B, Daubin V, Anthouard V, Aiach N, Arnaiz O, Billaut A, Beisson J, Blanc I, Bouhouche K, Câmara F, Duharcourt S, Guigo R, Gogendeau D, Katinka M, et al (2006) Global trends of whole-genome duplications revealed by the ciliate Paramecium tetraurelia. Nature 444: 171–178

Awata J, Takada S, Standley C, Lechtreck KF, Bellvé KD, Pazour GJ, Fogarty KE & Witman GB (2014) NPHP4 controls ciliary trafficking of membrane proteins and large soluble proteins at the transition zone. J. Cell Sci. 127: 4714–4727

Barker AR, Renzaglia KS, Fry K & Dawe HR (2014) Bioinformatic analysis of ciliary transition zone proteins reveals insights into the evolution of ciliopathy networks. BMC Genomics 15: 531

Basiri ML, Ha A, Chadha A, Clark NM, Polyanovsky A, Cook B & Avidor-Reiss T (2014) A migrating ciliary gate compartmentalizes the site of axoneme assembly in Drosophila spermatids. Curr. Biol. CB 24: 2622–2631

Blum JJ (1971) Existence of a breaking point in cilia and flagella. J. Theor. Biol. 33: 257–263

Boisvieux-Ulrich E, Sandoz D & Chailley B (1980) A thin and freeze-fracture study of deciliation in bird oviduct. Biol. Cell. 37: 261–268

Brenner RM (1969) Renewal of oviduct cilia during the menstrual cycle of the rhesus monkey. Fertil. Steril. 20: 599–611

Callen AM, Adoutte A, Andrew JM, Baroin-Tourancheau A, Bré MH, Ruiz PC, Clérot JC, Delgado P, Fleury A & Jeanmaire-Wolf R (1994) Isolation and characterization of libraries of monoclonal antibodies directed against various forms of tubulin in Paramecium. Biol. Cell 81: 95–119

Carradec Q, Götz U, Arnaiz O, Pouch J, Simon M, Meyer E & Marker S (2015) Primary and secondary siRNA synthesis triggered by RNAs from food bacteria in the ciliate Paramecium tetraurelia. Nucleic Acids Res. 43: 1818–1833

Carson JL, Collier AM & Hu SC (1980) Ultrastructural observations on cellular and subcellular aspects of experimental Mycoplasma pneumoniae disease. Infect. Immun. 29: 1117–1124

Chávez M, Ena S, Van Sande J, de Kerchove d’Exaerde A, Schurmans S & Schiffmann SN (2015) Modulation of Ciliary Phosphoinositide Content Regulates Trafficking and Sonic Hedgehog Signaling Output. Dev. Cell 34: 338–350

Craige B, Tsao C-C, Diener DR, Hou Y, Lechtreck K-F, Rosenbaum JL & Witman GB (2010) CEP290 tethers flagellar transition zone microtubules to the membrane and regulates flagellar protein content. J. Cell Biol. 190: 927–940

Czarnecki PG & Shah JV (2012) The ciliary transition zone: from morphology and molecules to medicine. Trends Cell Biol. 22: 201–210

Das RM & Storey KG (2014) Apical abscission alters cell polarity and dismantles the primary cilium during neurogenesis. Science 343: 200–204

Dean S, Moreira-Leite F, Varga V & Gull K (2016) Cilium transition zone proteome reveals compartmentalization and differential dynamics of ciliopathy complexes. Proc. Natl. Acad. Sci. U. S. A. 113: E5135–5143

Diener DR, Lupetti P & Rosenbaum JL (2015) Proteomic analysis of isolated ciliary transition zones reveals the presence of ESCRT proteins. Curr. Biol. CB 25: 379–384

Donnez J, Casanas-Roux F, Caprasse J, Ferin J & Thomas K (1985) Cyclic changes in ciliation, cell height, and mitotic activity in human tubal epithelium during reproductive life. Fertil. Steril. 43: 554–559

Dute R & Kung C (1978) Ultrastructure of the proximal region of somatic cilia in Paramecium tetraurelia. J. Cell Biol. 78: 451–464

Fassad MR, Shoemark A, le Borgne P, Koll F, Patel M, Dixon M, Hayward J, Richardson C, Frost E, Jenkins L, Cullup T, Chung EMK, Lemullois M, Aubusson-Fleury A, Hogg C, Mitchell DR, Tassin A-M & Mitchison HM (2018a) C11orf70 Mutations Disrupting the Intraflagellar Transport-Dependent Assembly of Multiple Axonemal Dyneins Cause Primary Ciliary Dyskinesia. Am. J. Hum. Genet. 102: 956–972

Fassad MR, Shoemark A, Legendre M, Hirst RA, Koll F, le Borgne P, Louis B, Daudvohra F, Patel MP, Thomas L, Dixon M, Burgoyne T, Hayes J, Nicholson AG, Cullup T, Jenkins L, Carr SB, Aurora P, Lemullois M, Aubusson-Fleury A, et al (2018b) Mutations in Outer Dynein Arm Heavy Chain DNAH9 Cause Motile Cilia Defects and Situs Inversus. Am. J. Hum. Genet. 103: 984–994

Finst RJ, Kim PJ, Griffis ER & Quarmby LM (2000) Fa1p is a 171 kDa protein essential for axonemal microtubule severing in Chlamydomonas. J. Cell Sci. 113 (Pt 11): 1963–1971

Finst RJ, Kim PJ & Quarmby LM (1998) Genetics of the deflagellation pathway in Chlamydomonas. Genetics 149: 927–936

Galvani A & Sperling L (2002) RNA interference by feeding in Paramecium. Trends Genet. TIG 18: 11–12

Garcia-Gonzalo FR, Corbit KC, Sirerol-Piquer MS, Ramaswami G, Otto EA, Noriega TR, Seol AD, Robinson JF, Bennett CL, Josifova DJ, García-Verdugo JM, Katsanis N, Hildebrandt F & Reiter JF (2011) A transition zone complex regulates mammalian ciliogenesis and ciliary membrane composition. Nat. Genet. 43: 776–784

Garcia-Gonzalo FR, Phua SC, Roberson EC, Garcia G, Abedin M, Schurmans S, Inoue T & Reiter JF (2015) Phosphoinositides Regulate Ciliary Protein Trafficking to Modulate Hedgehog Signaling. Dev. Cell 34: 400–409

Gibson DG, Young L, Chuang R-Y, Venter JC, Hutchison CA & Smith HO (2009) Enzymatic assembly of DNA molecules up to several hundred kilobases. Nat. Methods 6: 343–345

Gilley D, Preer JR, Aufderheide KJ & Polisky B (1988) Autonomous replication and addition of telomerelike sequences to DNA microinjected into Paramecium tetraurelia macronuclei. Mol. Cell. Biol. 8: 4765–4772

Goetz SC & Anderson KV (2010) The primary cilium: a signalling centre during vertebrate development. Nat. Rev. Genet. 11: 331–344

Gogendeau D, Guichard P & Tassin A-M (2015) Purification of centrosomes from mammalian cell lines. Methods Cell Biol. 129: 171–189

Gonçalves J & Pelletier L (2017) The Ciliary Transition Zone: Finding the Pieces and Assembling the Gate. Mol. Cells 40: 243–253

Heller RF & Gordon RE (1986) Chronic effects of nitrogen dioxide on cilia in hamster bronchioles. Exp. Lung Res. 10: 137–152

Hilton LK, Meili F, Buckoll PD, Rodriguez-Pike JC, Choutka CP, Kirschner JA, Warner F, Lethan M, Garces FA, Qi J & Quarmby LM (2016) A Forward Genetic Screen and Whole Genome Sequencing Identify Deflagellation Defective Mutants in Chlamydomonas, Including Assignment of ADF1 as a TRP Channel. G3 Bethesda Md 6: 3409–3418

Huang L, Szymanska K, Jensen VL, Janecke AR, Innes AM, Davis EE, Frosk P, Li C, Willer JR, Chodirker BN, Greenberg CR, McLeod DR, Bernier FP, Chudley AE, Müller T, Shboul M, Logan CV, Loucks CM, Beaulieu CL, Bowie RV, et al (2011) TMEM237 is mutated in individuals with a Joubert syndrome related disorder and expands the role of the TMEM family at the ciliary transition zone. Am. J. Hum. Genet. 89: 713–730

Iftode F, Cohen J, Ruiz F, Torres-Rueda A, Chen-Sban L, Adoutte A & Beisson J (1989) Development of surface pattern during division in Paramecium. I. Mapping of duplication and reorganization of cortical cytoskeletal structures in the wild type. development 105: 191–211

Jana SC, Mendonça S, Machado P, Werner S, Rocha J, Pereira A, Maiato H & Bettencourt-Dias M (2018) Differential regulation of transition zone and centriole proteins contributes to ciliary base diversity. Nat. Cell Biol. 20: 928–941

Janke C & Bulinski JC (2011) Post-translational regulation of the microtubule cytoskeleton: mechanisms and functions. Nat. Rev. Mol. Cell Biol. 12: 773–786

Laligné C, Klotz C, de Loubresse NG, Lemullois M, Hori M, Laurent FX, Papon JF, Louis B, Cohen J & Koll F (2010) Bug22p, a conserved centrosomal/ciliary protein also present in higher plants, is required for an effective ciliary stroke in Paramecium. Eukaryot. Cell 9: 645–655

Lambacher NJ, Bruel A-L, van Dam TJP, Szymańska K, Slaats GG, Kuhns S, McManus GJ, Kennedy JE, Gaff K, Wu KM, van der Lee R, Burglen L, Doummar D, Rivière J-B, Faivre L, Attié-Bitach T, Saunier S, Curd A, Peckham M, Giles RH, et al (2016) TMEM107 recruits ciliopathy proteins to subdomains of the ciliary transition zone and causes Joubert syndrome. Nat. Cell Biol. 18: 122–131

Li C, Jensen VL, Park K, Kennedy J, Garcia-Gonzalo FR, Romani M, Mori RD, Bruel A-L, Gaillard D, Doray B, Lopez E, Rivière J-B, Faivre L, Thauvin-Robinet C, Reiter JF, Blacque OE, Valente EM & Leroux MR (2016) MKS5 and CEP290 Dependent Assembly Pathway of the Ciliary Transition Zone. PLOS Biol. 14: e1002416

Lohret TA, McNally FJ & Quarmby LM (1998) A role for katanin-mediated axonemal severing during Chlamydomonas deflagellation. Mol. Biol. Cell 9: 1195–1207

Lohret TA, Zhao L & Quarmby LM (1999) Cloning of Chlamydomonas p60 katanin and localization to the site of outer doublet severing during deflagellation. Cell Motil. Cytoskeleton 43: 221–231

Mahjoub MR, Montpetit B, Zhao L, Finst RJ, Goh B, Kim AC & Quarmby LM (2002) The FA2 gene of Chlamydomonas encodes a NIMA family kinase with roles in cell cycle progression and microtubule severing during deflagellation. J. Cell Sci. 115: 1759–1768

Mirvis M, Siemers KA, Nelson WJ & Stearns T (2019) Primary Cilium Disassembly in Mammalian Cells Occurs Predominantly by Whole-Cilium Shedding. bioRxiv: 433144

Mitchison HM & Valente EM (2017) Motile and non-motile cilia in human pathology: from function to phenotypes. J. Pathol. 241: 294–309

Muse KE, Collier AM & Baseman JB (1977) Scanning electron microscopic study of hamster tracheal organ cultures infected with Bordetella pertussis. J. Infect. Dis. 136: 768–777

Nelson DL (1995) Chapter 4 Preparation of Cilia and Subciliary Fractions from Paramecium. In Methods in Cell Biology, Dentler W & Witman G (eds) pp 17–24. Academic Press Available at: http://www.sciencedirect.com/science/article/pii/S0091679X08607852 [Accessed June 18, 2019]

Odor DL, Gaddum-Rosse P, Rumery RE & Blandau RJ (1980) Cyclic variations in the oviuductal ciliated cells during the menstrual cycle and after estrogen treatment in the pig-tailed monkey, Macaca nemestrina. Anat. Rec. 198: 35–57

Ohno, S. Evolution by Gene Duplication (George Allen and Unwin, London, 1970).

Ott C, Nachmias D, Adar S, Jarnik M, Sherman S, Birnbaum RY, Lippincott-Schwartz J & Elia N (2018) VPS4 is a dynamic component of the centrosome that regulates centrosome localization of γ-tubulin, centriolar satellite stability and ciliogenesis. Sci. Rep. 8: 3353

Overgaard CE, Sanzone KM, Spiczka KS, Sheff DR, Sandra A & Yeaman C (2009) Deciliation is associated with dramatic remodeling of epithelial cell junctions and surface domains. Mol. Biol. Cell 20: 102–113

Papon JF, Perrault I, Coste A, Louis B, Gérard X, Hanein S, Fares-Taie L, Gerber S, Defoort-Dhellemmes S, Vojtek AM, Kaplan J, Rozet JM & Escudier E (2010) Abnormal respiratory cilia in non-syndromic Leber congenital amaurosis with CEP290 mutations. J. Med. Genet. 47: 829–834

Perrone CA, Myster SH, Bower R, O’Toole ET & Porter ME (2000) Insights into the structural organization of the I1 inner arm dynein from a domain analysis of the 1beta dynein heavy chain. Mol. Biol. Cell 11: 2297–2313

Phua SC, Chiba S, Suzuki M, Su E, Roberson EC, Pusapati GV, Setou M, Rohatgi R, Reiter JF, Ikegami K & Inoue T (2017) Dynamic Remodeling of Membrane Composition Drives Cell Cycle through Primary Cilia Excision. Cell 168: 264–279.e15

Praetorius HA & Spring KR (2003) Removal of the MDCK cell primary cilium abolishes flow sensing. J. Membr. Biol. 191: 69–76

Quarmby LM (1996) Ca2+ influx activated by low pH in Chlamydomonas. J. Gen. Physiol. 108: 351–361

Quarmby LM (2004) Cellular deflagellation. Int. Rev. Cytol. 233: 47–91

Quarmby LM (2009) Chapter 3 - Deflagellation. In The Chlamydomonas Sourcebook (Second Edition), Harris EH, Stern DB & Witman GB (eds) pp 43–69. London: Academic Press Available at: http://www.sciencedirect.com/science/article/pii/B978012370873100040X [Accessed May 27, 2019]

Quarmby LM & Hartzell HC (1994) Two distinct, calcium-mediated, signal transduction pathways can trigger deflagellation in Chlamydomonas reinhardtii. J. Cell Biol. 124: 807–815

Quarmby LM, Yueh YG, Cheshire JL, Keller LR, Snell WJ & Crain RC (1992) Inositol phospholipid metabolism may trigger flagellar excision in Chlamydomonas reinhardtii. J. Cell Biol. 116: 737–744

Reiter JF, Blacque OE & Leroux MR (2012) The base of the cilium: roles for transition fibres and the transition zone in ciliary formation, maintenance and compartmentalization. EMBO Rep. 13: 608–618

Rüdiger AH, Rüdiger M, Wehland J & Weber K (1999) Monoclonal antibody ID5: epitope characterization and minimal requirements for the recognition of polyglutamylated alpha- and beta-tubulin. Eur. J. Cell Biol. 78: 15–20

Rumery RE & Eddy EM (1974) Scanning electron microscopy of the fimbriae and ampullae of rabbit oviducts. Anat. Rec. 178: 83–101

Sang L, Miller JJ, Corbit KC, Giles RH, Brauer MJ, Otto EA, Baye LM, Wen X, Scales SJ, Kwong M, Huntzicker EG, Sfakianos MK, Sandoval W, Bazan JF, Kulkarni P, Garcia-Gonzalo FR, Seol AD, O’Toole JF, Held S, Reutter HM, et al (2011) Mapping the NPHP-JBTS-MKS protein network reveals ciliopathy disease genes and pathways. Cell 145: 513–528

Satir B, Sale WS & Satir P (1976) Membrane renewal after dibucaine deciliation of Tetrahymena. Freeze-fracture technique, cilia, membrane structure. Exp. Cell Res. 97: 83–91

Schouteden C, Serwas D, Palfy M & Dammermann A (2015) The ciliary transition zone functions in cell adhesion but is dispensable for axoneme assembly in C. elegans. J. Cell Biol. 210: 35–44

Shi L, Koll F, Arnaiz O & Cohen J (2018) The Ciliary Protein IFT57 in the Macronucleus of Paramecium. J. Eukaryot. Microbiol. 65: 12–27

Skouri F & Cohen J (1997) Genetic approach to regulated exocytosis using functional complementation in Paramecium: identification of the ND7 gene required for membrane fusion. Mol. Biol. Cell 8: 1063–1071

Sonneborn TM (1970) Chapter 12 Methods in Paramecium Research**With contributed sections by: S. Dryl, Nencki Institute of Experimental Biology, Warsaw, Poland; E. D. Hanson, Connecticut Wesleyan University, Middletown, Connecticut; K. Hiwatashi, Tôhoku University, Sendai, Japan; S. Koizumi, University of Iwate, Ueda Morioka, Japan; A. Miyake, Kyoto University, Kyoto, Japan; J. R. Preer, Jr., Indiana University, Bloomington, Indiana; A. Reisner, CSIRO, division of Animal Genetics, Epping, New South Wales; E. Steers, Department of Health, Education and Welfare, National Institute of Health, Bethesda, Maryland; S. Taub, Princeton University, Princeton, New Jersey. In Methods in Cell Biology, Prescott DM (ed) pp 241–339. Academic Press Available at: http://www.sciencedirect.com/science/article/pii/S0091679X08617586 [Accessed May 28, 2019]

Suchard SJ, Rhoads DE & Kaneshiro ES (1989) The inositol lipids of Paramecium tetraurelia and preliminary characterizations of phosphoinositide kinase activity in the ciliary membrane. J. Protozool. 36: 185–190

Tassin A-M, Lemullois M & Aubusson-Fleury A (2015) Paramecium tetraurelia basal body structure. Cilia 5: 6

Timmons L & Fire A (1998) Specific interference by ingested dsRNA. Nature 395: 854

Verhage HG, Bareither ML, Jaffe RC & Akbar M (1979) Cyclic changes in ciliation, secretion and cell height of the oviductal epithelium in women. Am. J. Anat. 156: 505–521

Wehland J & Weber K (1987) Turnover of the carboxy-terminal tyrosine of alpha-tubulin and means of reaching elevated levels of detyrosination in living cells. J. Cell Sci. 88 (Pt 2): 185–203

Weng RR, Yang TT, Huang C-E, Chang C-W, Wang W-J & Liao J-C (2018) Super-Resolution Imaging Reveals TCTN2 Depletion-Induced IFT88 Lumen Leakage and Ciliary Weakening. Biophys. J. 115: 263–275

Wheeler GL, Joint I & Brownlee C (2008) Rapid spatiotemporal patterning of cytosolic Ca2+ underlies flagellar excision in Chlamydomonas reinhardtii. Plant J. Cell Mol. Biol. 53: 401–413

Wiegering A, Dildrop R, Kalfhues L, Spychala A, Kuschel S, Lier JM, Zobel T, Dahmen S, Leu T, Struchtrup A, Legendre F, Vesque C, Schneider-Maunoury S, Saunier S, Rüther U & Gerhardt C (2018a) Cell type-specific regulation of ciliary transition zone assembly in vertebrates. EMBO J. 37:

Wiegering A, Rüther U & Gerhardt C (2018b) The ciliary protein Rpgrip1l in development and disease. Dev. Biol. 442: 60–68

Williams CL, Li C, Kida K, Inglis PN, Mohan S, Semenec L, Bialas NJ, Stupay RM, Chen N, Blacque OE, Yoder BK & Leroux MR (2011) MKS and NPHP modules cooperate to establish basal body/transition zone membrane associations and ciliary gate function during ciliogenesis. J. Cell Biol. 192: 1023–1041

Wilson R, Read R, Thomas M, Rutman A, Harrison K, Lund V, Cookson B, Goldman W, Lambert H & Cole P (1991) Effects of Bordetella pertussis infection on human respiratory epithelium in vivo and in vitro. Infect. Immun. 59: 337–345

Yang TT, Su J, Wang W-J, Craige B, Witman GB, Tsou M-FB & Liao J-C (2015) Superresolution Pattern Recognition Reveals the Architectural Map of the Ciliary Transition Zone. Sci. Rep. 5: 14096

Yano J, Rajendran A, Valentine MS, Saha M, Ballif BA & Van Houten JL (2013) Proteomic analysis of the cilia membrane of Paramecium tetraurelia. J. Proteomics 78: 113–122

Yueh YG & Crain RC (1993) Deflagellation of Chlamydomonas reinhardtii follows a rapid transitory accumulation of inositol 1,4,5-trisphosphate and requires Ca2+ entry. J. Cell Biol. 123: 869–875

Zhang D & Aravind L (2012) Novel transglutaminase-like peptidase and C2 domains elucidate the structure, biogenesis and evolution of the ciliary compartment. Cell Cycle Georget. Tex 11: 3861–3875

