## Supplemental Table1 for "MKS-NPHP module proteins regulate ciliary shedding in *Paramecium*"

**Table S1: ID list of transition zone proteins in *Paramecium***

| GENE NAME | GENE ID |
|---|---|
| MKS1 | PTET.51.1.G0320263 |
| MKS2/TMEM216 | PTET.51.1.G0150351<br>PTETP2500008001<br>PTET.51.1.G0300269 |
| MKS3/TMEM67/MECKELIN | PTET.51.1.G0460260 |
| AHI1 | PTET.51.1.G0800017<br>PTET.51.1.G1620035 |
| B9D1 | PTET.51.1.G1150027<br>PTET.51.1.G1130142 |
| B9D2 | PTET.51.1.G1470056<br>PTET.51.1.G0610100<br>PTET.51.1.G0280188 |
| CC2D2A | PTET.51.1.G0190283 |
| TECTONIC (1,2 and 3) | PTET.51.1.G0700187 |
| TMEM17 | PTET.51.1.P0500202<br>PTET.51.1.P1210132<br>PTET.51.1.P0380214 |
| TMEM107 | PTET.51.1.G0300250<br>PTET.51.1.P0310017 |
| TMEM218 | - |
| TMEM231 | PTET.51.1.P1340134<br>PTET.51.1.P0010036 |
| TMEM237 | - |
| NPHP1 | PTET.51.1.G5560037<br>PTET.51.1.G1530007 |
| NPHP4/POC10 | PTET.51.1.P0020231<br>PTET.51.1.G0220034<br>PTET.51.1.P0290127<br>PTET.51.1.P0130026 |
| NPHP5/IQCB1 | - |
| NPHP3/MKS7 | PTET.51.1.P0280102<br>PTET.51.1.P0610178<br>PTET.51.1.G0340120<br>PTET.51.1.P0480171 |
| CEP290/NPHP6/MKS4 | PTET.51.1.G0130345<br>PTET.51.1.P0190305 |
| RPGRIP1L/NPHP8 | PTET.51.1.G0340252<br>PTET.51.1.G0480042 |
