## Supplementary figures and images for "MKS-NPHP module proteins regulate ciliary shedding in *Paramecium*"

### Supplemental Figure 1

scale=10 μm
scale=1 μm

GOGENDEAU-et-al

#### FIGURE S1: CILIATION STATUS IN *PARAMECIUM*

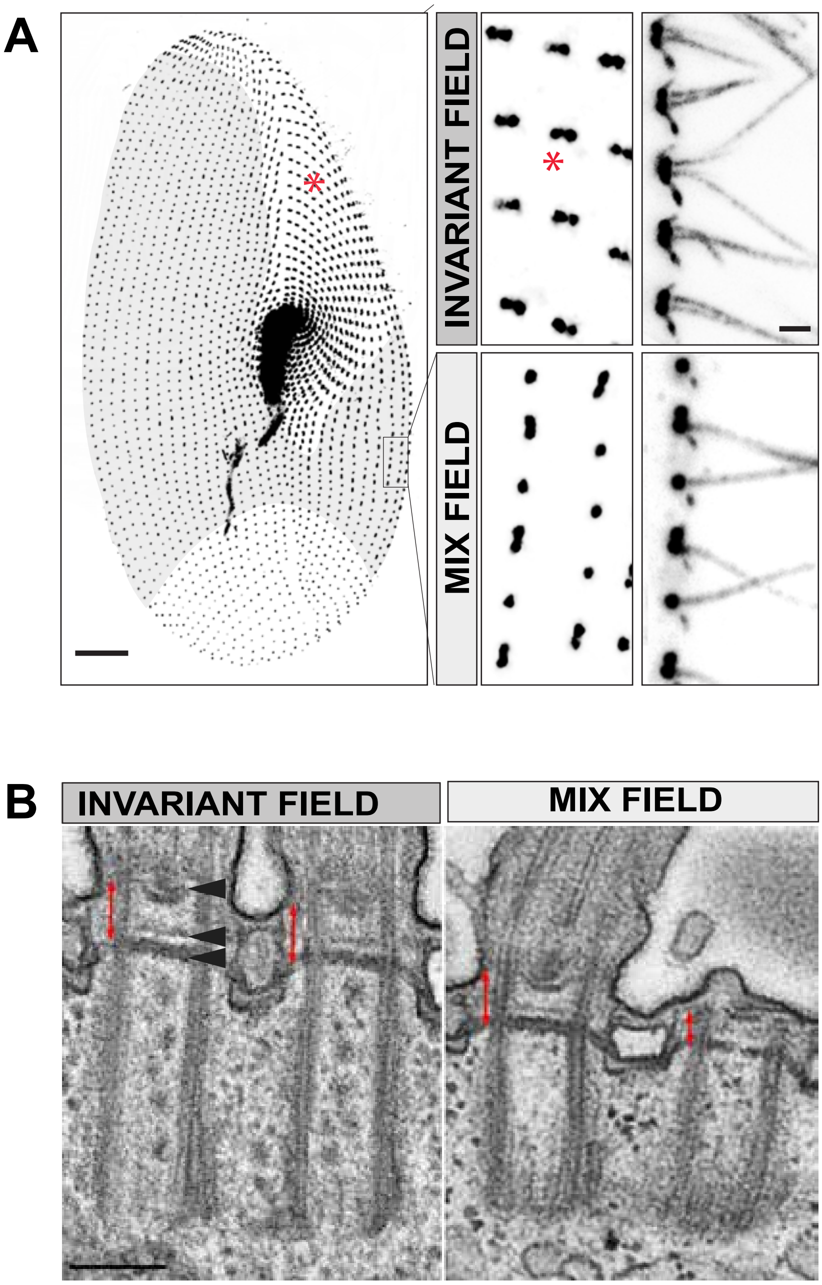

### Supplemental Figure 2

GOGENDEAU-et-al

FIGURE S2: PARAMECIUM TRANSITION ZONE PROTEINS ARE RECRUITED WHEN CILIA EXTEND

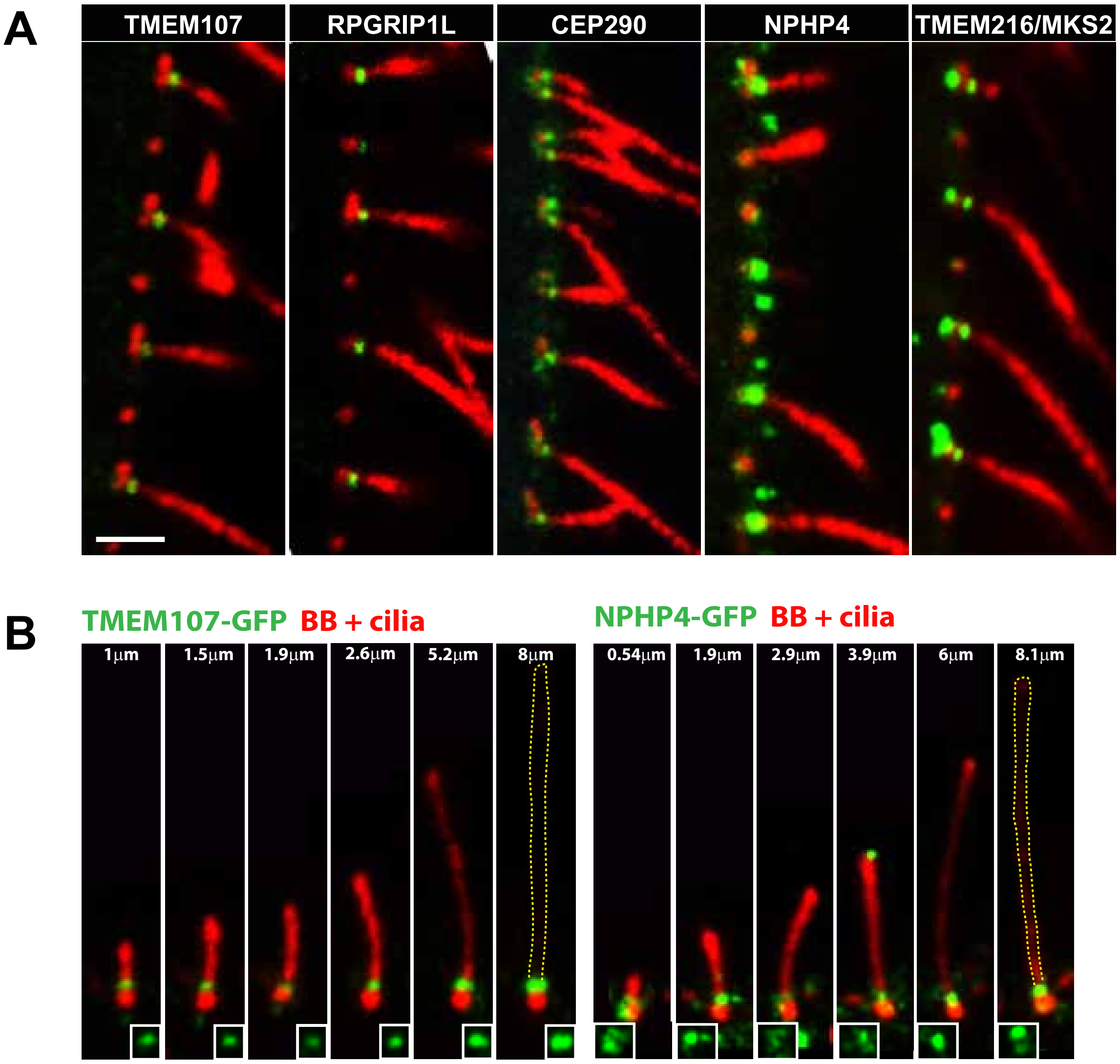

### Supplemental Figure 3

#### GOGENDEAU-et-al
#### FIGURE S3: Efficiency of inactivation of the different RNAi vectors

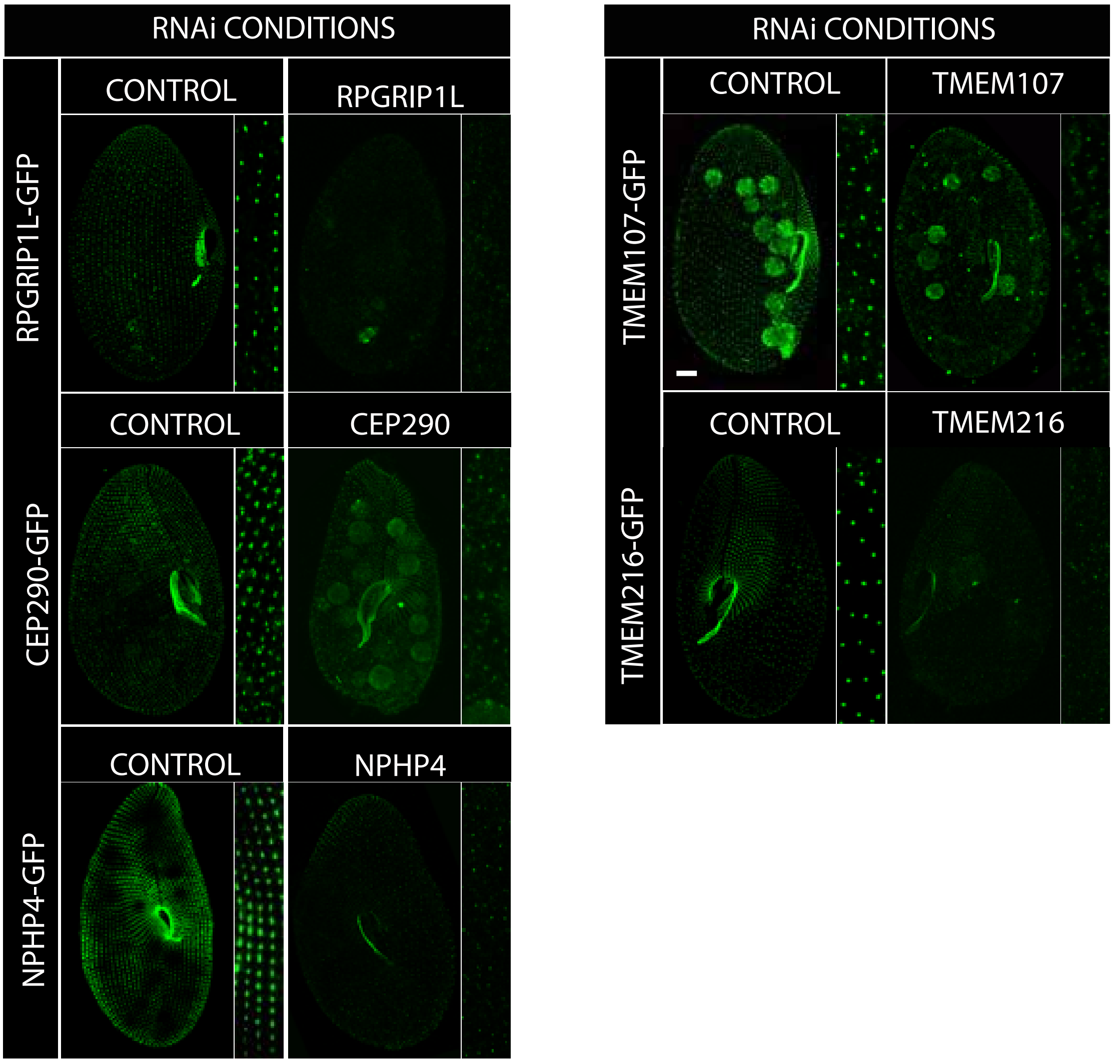

### Supplemental Figure 4

#### GOGENDEAU-et-al

#### FIGURE S4: DEPLETION OF TZ PROTEINS AFFECTS SWIMMING BEHAVIOR

A

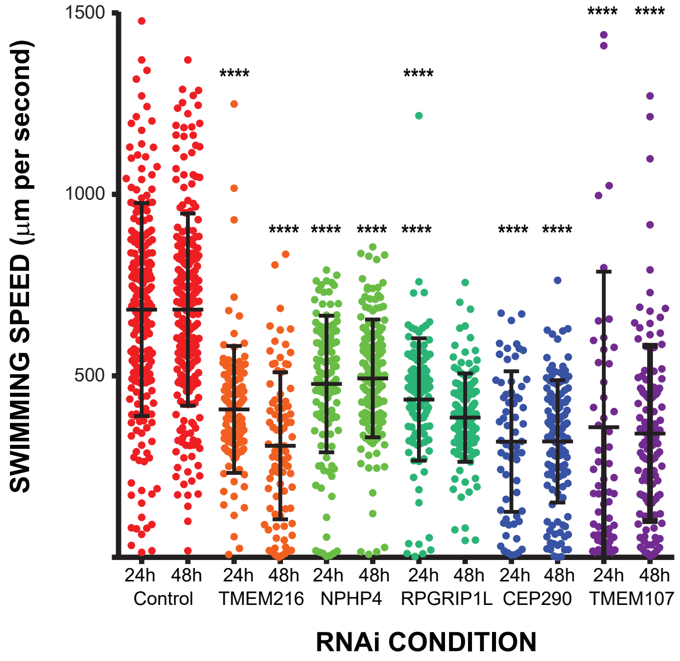

### Supplemental Figure 8

GOGENDEAU-et-al

FIGURE S8: RPGRIP1L CONSERVED C DOMAINS ARE INVOLVED IN THE DECILIATION SIGNAL

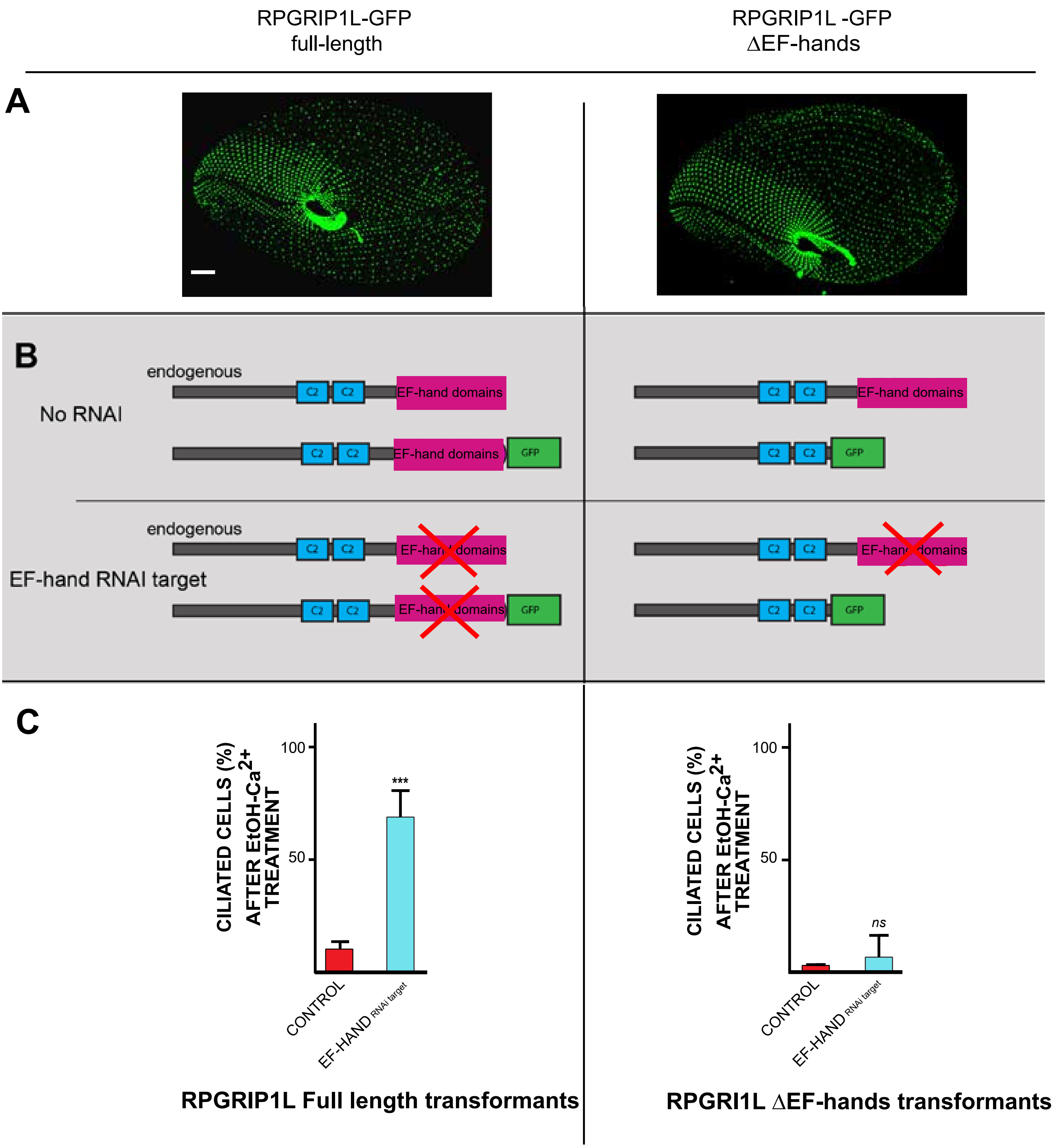
