## Supplemental Figure 5 for "MKS-NPHP module proteins regulate ciliary shedding in *Paramecium*"

GOGENDEAU-et-al

### SUPPLEMENTAL FIGURE 5: DEPLETION OF TZ PROTEINS DO NOT AFFECT BASAL BODY POSITIONING

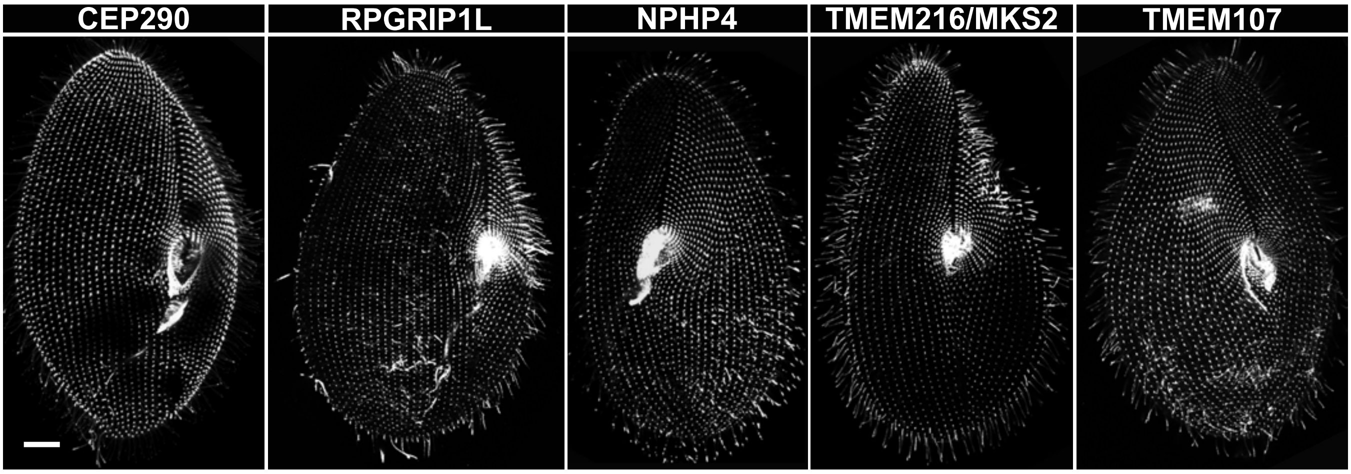
