## Supplemental Figure 6 for "MKS-NPHP module proteins regulate ciliary shedding in *Paramecium*"

GOGENDEAU-et-al

### SUPPLEMENTAL FIGURE 6: TMEM107 AND TMEM216 DEPLETED CELLS SHED THEIR CILIA DISTALLY OF THE TRANSITION ZONE

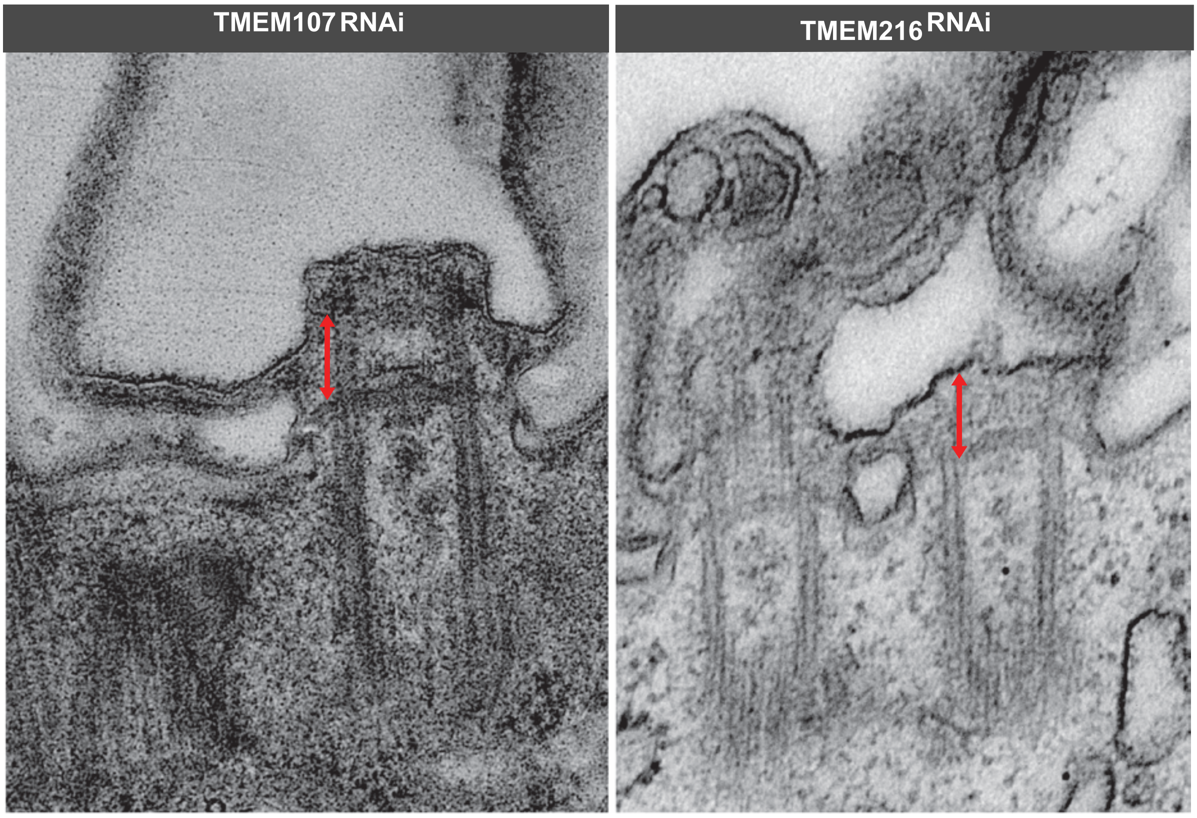
