## Supplemental Figure 7 for "MKS-NPHP module proteins regulate ciliary shedding in *Paramecium*"

**GOGENDEAU-et-al**

**SUPPLEMENTAL FIGURE 7: Heatmaps corresponding to transcriptomic analyses of TMEM216$^{RNAi}$ compared to control and IFT 57$^{RNAi}$**

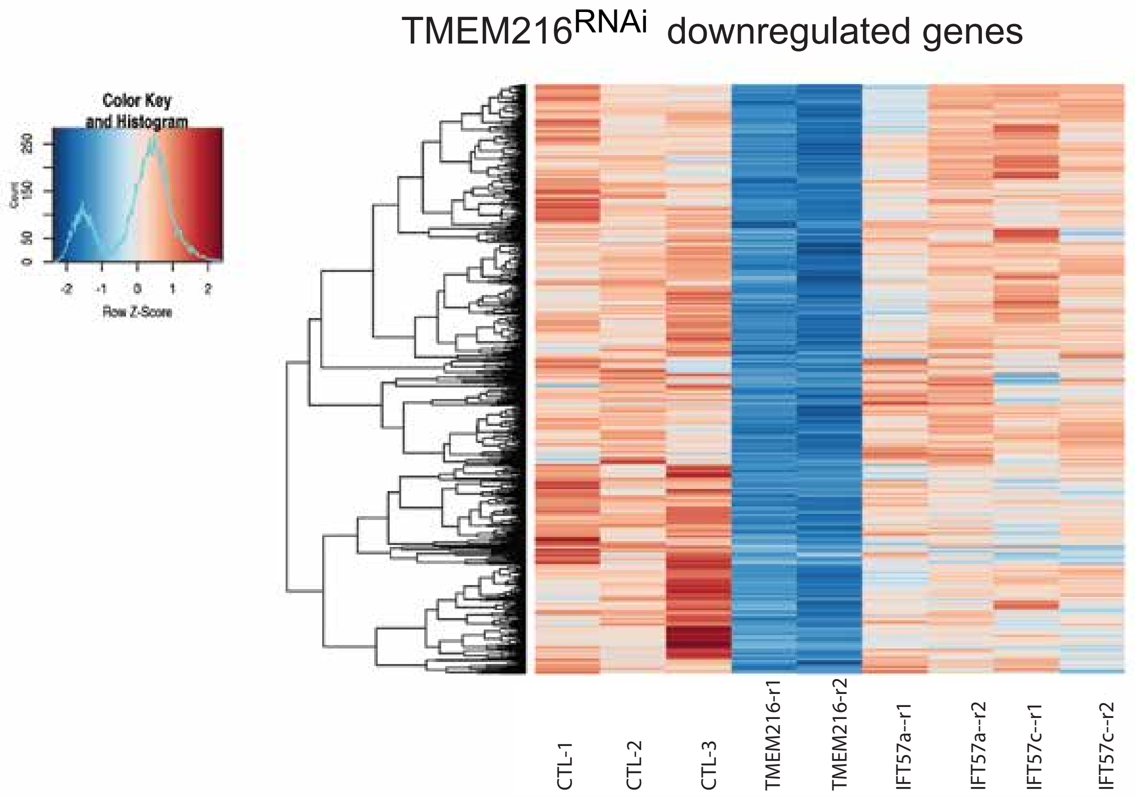


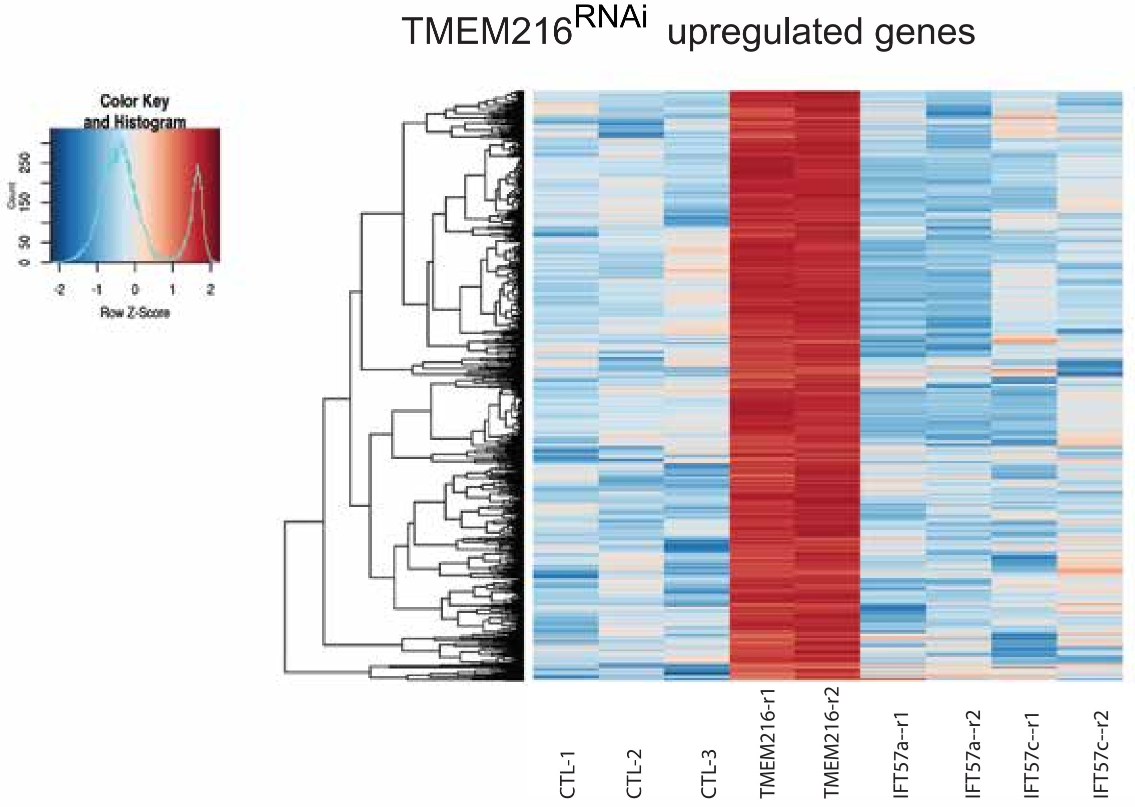
